# Structural basis for lysophosphatidylserine recognition by GPR34

**DOI:** 10.1101/2023.02.15.528751

**Authors:** Tamaki Izume, Ryo Kawahara, Akiharu Uwamizu, Luying Chen, Shun Yaginuma, Jumpei Omi, Hiroki Kawana, Fumiya K. Sano, Tatsuki Tanaka, Kazuhiro Kobayashi, Hiroyuki H. Okamoto, Yoshiaki Kise, Tomohiko Ohwada, Junken Aoki, Wataru Shihoya, Osamu Nureki

## Abstract

GPR34 is a recently identified G-protein coupled receptor, which has an immunomodulatory role and recognizes lysophosphatidylserine (LysoPS) as a putative ligand. Here, we report cryo-electron microscopy structures of human GPR34-G_i_ complex bound with either the LysoPS analogue S3E-LysoPS, which contains an ethoxy group at the *sn*-1 position, or M1, a derivative of S3E-LysoPS in which oleic acid is substituted with a metabolically stable aromatic fatty acid surrogate. In both structures, the ligand-binding pocket is laterally open toward the membrane, allowing lateral entry of lipidic agonists into the cavity. The amine and carboxylate groups of the serine moiety are recognized by the charged residue cluster, and the aromatic fatty acid surrogate of M1 forms stable hydrophobic interactions with the cavity, thus acting as a superagonist. Molecular dynamics simulations further account for the LysoPS-regioselectivity of GPR34. Thus, using a series of structural and physiological experiments, we provide evidence that chemically unstable 2-acyl LysoPS is the physiological ligand for GPR34, suggesting its short signal duration. Overall, we anticipate the present structures will pave the way for development of novel anticancer drugs that specifically target GPR34.

## Main

GPR34 is a G-protein coupled receptor (GPCR) that is evolutionarily conserved in vertebrates^1–3^ and shows a high degree of homology to P2Y family members. In various species, GPR34 is expressed in a wide range of tissues and cells, including immune cells, such as microglia^4,5^, macrophages^6^, type 3 innate lymphoid cells^7^, platelets, and dendritic cells^8^. Previous studies have indicated that GPR34 is involved in numerous processes, which include the repair of damaged tissues by type 3 innate lymphoid cells^7^, activation of microglial phagocytosis^9^, neuropathy pain onset^10^, dendritic cell survival^8^, and suppression of infection^11^. Despite these myriad functions, the essential roles of GPR34 remain to be elucidated, primarily due to a lack of consensus regarding the identity of endogenous GPR34 ligands. Two previous studies by Sugo *et al*^12^ and Inoue *et al*^13^. identified lysophosphatidylserine (LysoPS) as the GPR34 ligand. This finding prompted Inoue *et al*^13^. to propose renaming GPR34 as LPS_1_ or LPSR1, based on its reported ability to recognize lysophosphatidic acid (LPA_1–6_). Other such LysoPS receptors, including P2Y10 (LPS_2_) and GPR174 (LPS_3_), have also been identified. However, the question of whether LysoPS is truly the physiological ligand for GPR34 remains controversial. In particular, Liebscher and colleagues were unable to replicate the finding by Sugo *et al*^12^. that LysoPS activates mouse and human GPR34 in cAMP inhibition assays^11^. Interestingly, however, the same group showed that two GPR34s from carp fish nicely respond to LysoPS in the same assay. Thus, further work is needed to confirm the identity of the physiological ligand for GPR34, and one possible approach is via the structural determination of the GPR34–LysoPS complex. LysoPS consists of L-serine and fatty acid moieties connected to a central glycerol molecule by phosphodiester and ester linkages. Our previous ligand structure–function relationship (SAR) studies using chemically modified LysoPS have demonstrated that both the serine and lipid moieties are required for GPR34 activation^14–16^. Physiologically, LysoPS is generated when phosphatidylserine-specific phospholipase A1 (PS-PLA1) hydrolyses PS regio-selectively at the *sn*-1 position to produce *sn*-2 LysoPS (Fig. 1a); *sn*-1 LysoPS is then easily formed by non-enzymatic migration of the ester^17^. Thus, both *sn*-1 and *sn*-2 LysoPS are found in mammals and are biologically active. Notably, GPR34 regio-selectively prefers LysoPS with an unsaturated fatty acid at the *sn*-2 position^1^. Such regioselectivity is not observed in other lipid-sensing GPCRs. Indeed, we previously performed SAR studies with synthetic LysoPS analogues that mimic *sn*-1 and *sn*-2 LysoPS and, in doing so, identified GPR34-, P2Y10-, and GPR174-selective agonists. Moreover, although the *in vivo* existence and biological activities of *sn*-3 lysophospholipids remain enigmatic, we synthesized LysoPS analogues with the *sn*-3 configuration and found that these show high potency and selectivity for GPR34^18^. In one case, by replacing the fatty acid with an aromatic group, we developed a metabolically stable GPR34 agonist, named M1^18^ (Fig. 1a). Critically, such *sn*-3 LysoPS derivatives represent valuable tools and may hold potential as therapeutic agonists of GPR34.

**Fig. 1:**
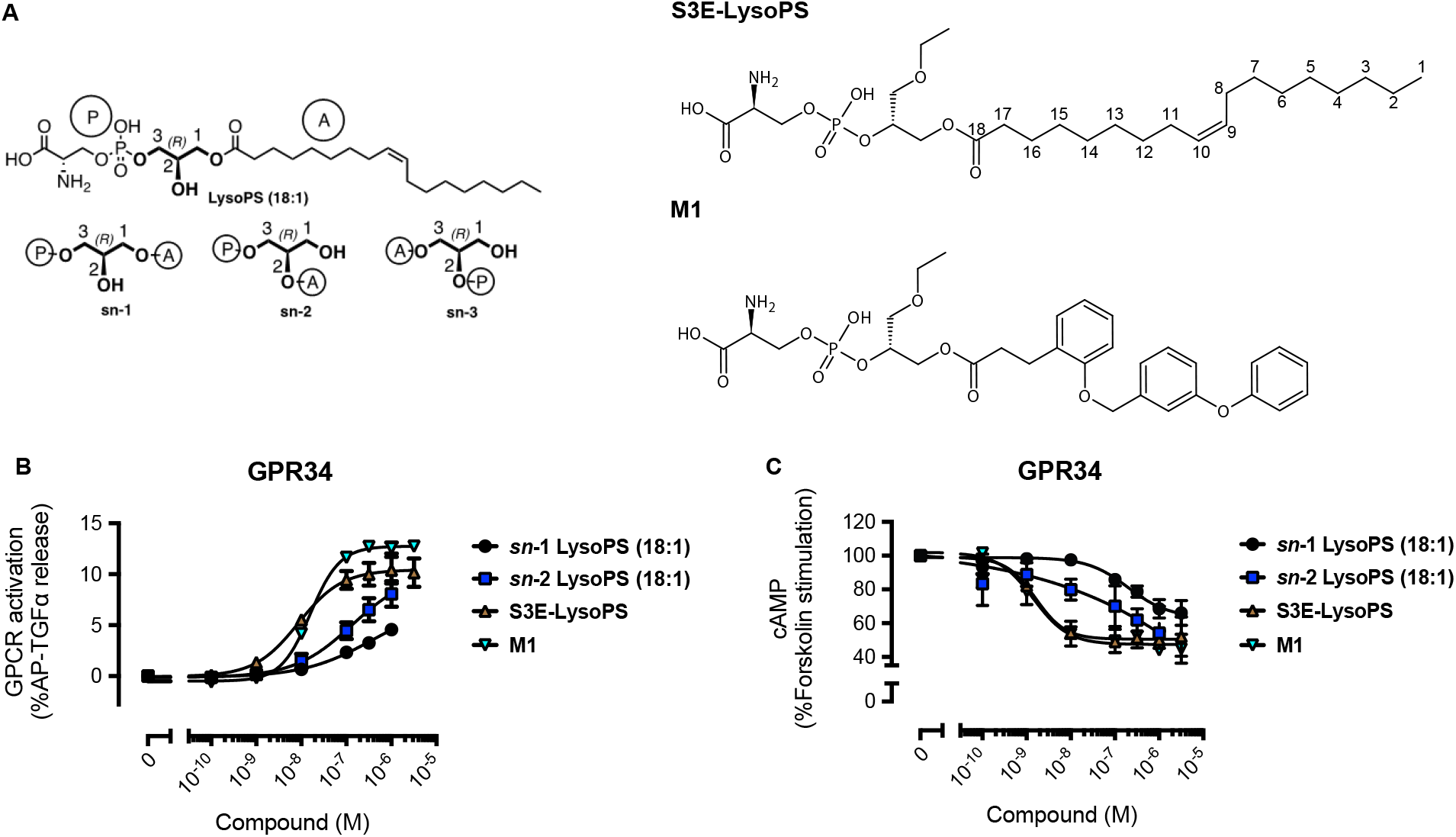
GPR34/LPS1 functions to potentiate immune responses *in vivo*. (**a**) Chemical structures of the GPR34 agonists used in this study. (**b and c**) Agonist activities of each lysophosphatidylserine (LysoPS) molecule toward GPR34, measured by the transforming growth factor (TGF)-α shedding assay **(b)** and the cyclic adenosine monophosphate (cAMP) assay **(c)**. Both *sn*-1 LysoPS (18:1) and *sn*-2 LysoPS (18:1) were prepared by digesting di-oleoyl (18:1) PS with *Rhizomucor miehei* lipase, which has phospholipase A1 activity. **(b)** For the TGF-α shedding assay, HEK293A cells expressing alkaline phosphatase-tagged (AP)-TGFα, Gαq/i1, and GPR34, as well as negative-control cells transfected with empty plasmid in place of the GPR34-encoding plasmid, were treated with each LysoPS. After 1 h incubation, the amount of AP-TGFα in the supernatant was determined. Receptor-specific AP-TGFα release, measured as the difference in AP-TGFα release between GPR34-expressing and negative control cells, is shown for each agonist. **(c)** In the cAMP assay, HEK293 cells expressing GPR34 and GloSensor-22F, a cAMP biosensor, were loaded with D-Luciferin and treated with each LysoPS plus forskolin. Luminescence signals were then measured and normalized, and the ratios of cAMP signal when stimulated with LysoPS *vs*. forskolin only are shown. Symbols and error bars indicate the average and standard error of the mean (SEM), respectively, from three independent experiments.\

Previous reports have suggested an immunomodulatory role for GPR34 signalling^7,9,11^; however, as noted above, it remains elusive whether LysoPS is a genuine *in vivo* ligand for GPR34, or how LysoPS activates GPR34 at the molecular level, limiting the drug development of the GPR34-targeting strategy. Here, we report two cryo-electron microscopy (cryo-EM) structures of human GPR34-G_i_ complex bound to an *sn*-3 LysoPS derivative and the potent M1 agonist. These structures, combined with results from molecular dynamics (MD) simulations, reveal the regioselectivity of LysoPS, the lipid-mimetic binding of stable agonist, and the unique G-protein coupling mode.

## Results

### GPR34 agonist ligands

GPR34 agonists used in this study are shown in Figure 1a. Depending on the fatty acid position in the glycerol backbone, LysoPS molecules are classified into one of three types (*sn*-1, *sn*-2 or *sn*-3)^18^. S3E-LysoPS is an analogue of *sn*-3 18:1 LysoPS, with an ethoxy group at the *sn*-1 position. M1 is an analogue of S3E-LysoPS, in which the fragile oleic acid is substituted with a more metabolically stable aromatic fatty acid surrogate (*i.e*., three tandemly linked phenyl groups, with two ether bonds)^18^. In transforming growth factor (TGF)-α shedding assays^13^ and cAMP assays, *sn*-2 LysoPS displays increased potency relative to *sn*-1 LysoPS, confirming that GPR34 prefers LysoPS with a fatty acid at the *sn*-2 position (Fig. 1b,c and Extended Data Tables 1 and 2). Notably, S3E-LysoPS and M1 yield EC_50_ values that are approximately 20-fold lower than those of *sn*-2 LysoPS, indicating increased potency for these agonists. Moreover, the E_max_ value for M1 is higher than for all other agonists, suggesting it acts as a superagonist.

### Overall structures

Full-length human GPR34 for cryo-EM analysis was expressed in HEK293 cells and purified with S3E-LysoPS or M1. Receptor was then incubated with the G_i_ heterotrimer (Gα_i1_, Gβ_1_, and Gγ_2_) and scFv16, which stabilizes GPCR-G_i_ complex formation, and the complex was purified by anti-Flag affinity and size exclusion chromatography. We then determined the structures of S3E-LysoPS- and M1-bound GPR34-G_i_ complexes at nominal global resolutions of 3.4 Å and 3.3 Å, respectively (Fig. 2a,b, Extended Data Fig. 1, and Extended Data Table 3). Local refinement with the mask of the receptor improved local resolution of the extracellular half of the receptor, and the resulting cryo-EM maps allowed modelling of the entire complexes, including agonists (Extended Data Fig. 2).

**Fig. 2:**
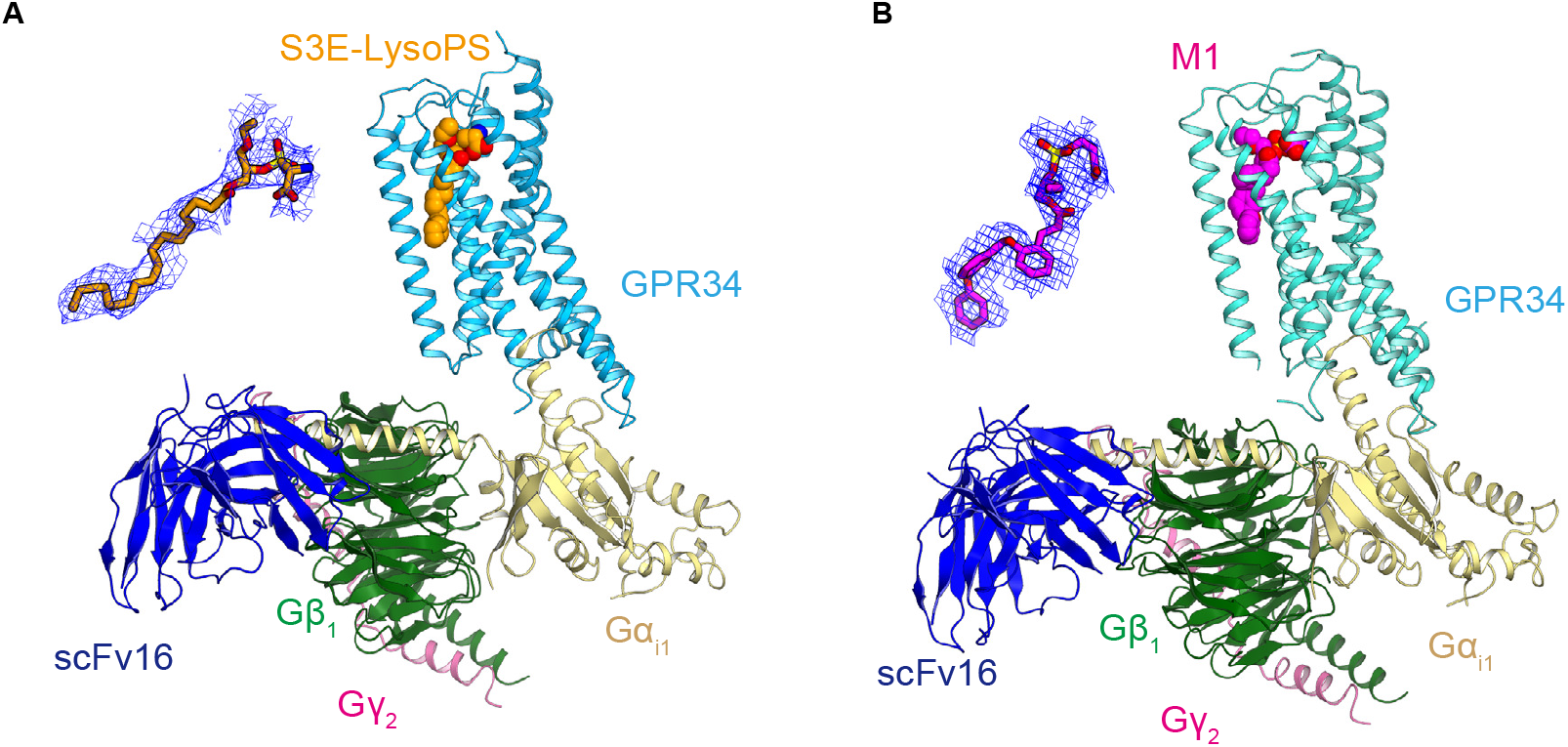
Cryo-electron microscopy (cryo-EM) structures of agonist-bound GPR34-G_i_. (**a and b**) Overall cryo-EM structures of the **(a)** S3E-LysoPS- and **(b)** M1-bound GPR34-Gi complexes. The agonists are indicated by Corey–Pauling–Koltun (CPK) models, and densities of the agonists are also shown.

GPR34 adopts the canonical GPCR topology of a heptahelical transmembrane bundle (7TM), with an extracellular N-terminus, three extracellular loops (ECLs), three intracellular loops (ICLs), and a short amphipathic helix 8 (H8) oriented parallel to the membrane (Fig. 3a). In GPR34, the conserved P^5.50^ (superscripts indicate Ballesteros–Weinstein numbers^19^) is replaced by I230^5.50^, and thus, the transmembrane helix TM5 forms a straight helix. The N-terminus is anchored to TM7 by the disulfide bond C46^N-ter^–C299^7.25^ (Fig. 3b), which is typically observed in class A GPCRs^20,21^. ECL2 (residues 196–213) adopts a U-shape with the TM4–5 side open (Fig. 3b) and is anchored by the disulfide bond C127^3.25^–C204^ECL2^, which is highly conserved in class A GPCRs^22^. ECL2 fills the transmembrane pocket facing toward the extracellular side and provides extensive interactions with TM2–6 (Fig. 3b). Specifically, F205^ECL2^ protrudes into the pocket, and H206^ECL2^ and K210^ECL2^ form salt bridges with E50^1.28^ and E216^5.36^, respectively. The backbone carbonyl groups in ECL2 form hydrogen bonds with residues in TM2, 3, and 6. The tightly packed ECL2 limits the space within the transmembrane pocket and constitutes the ligand-binding site.

**Fig. 3:**
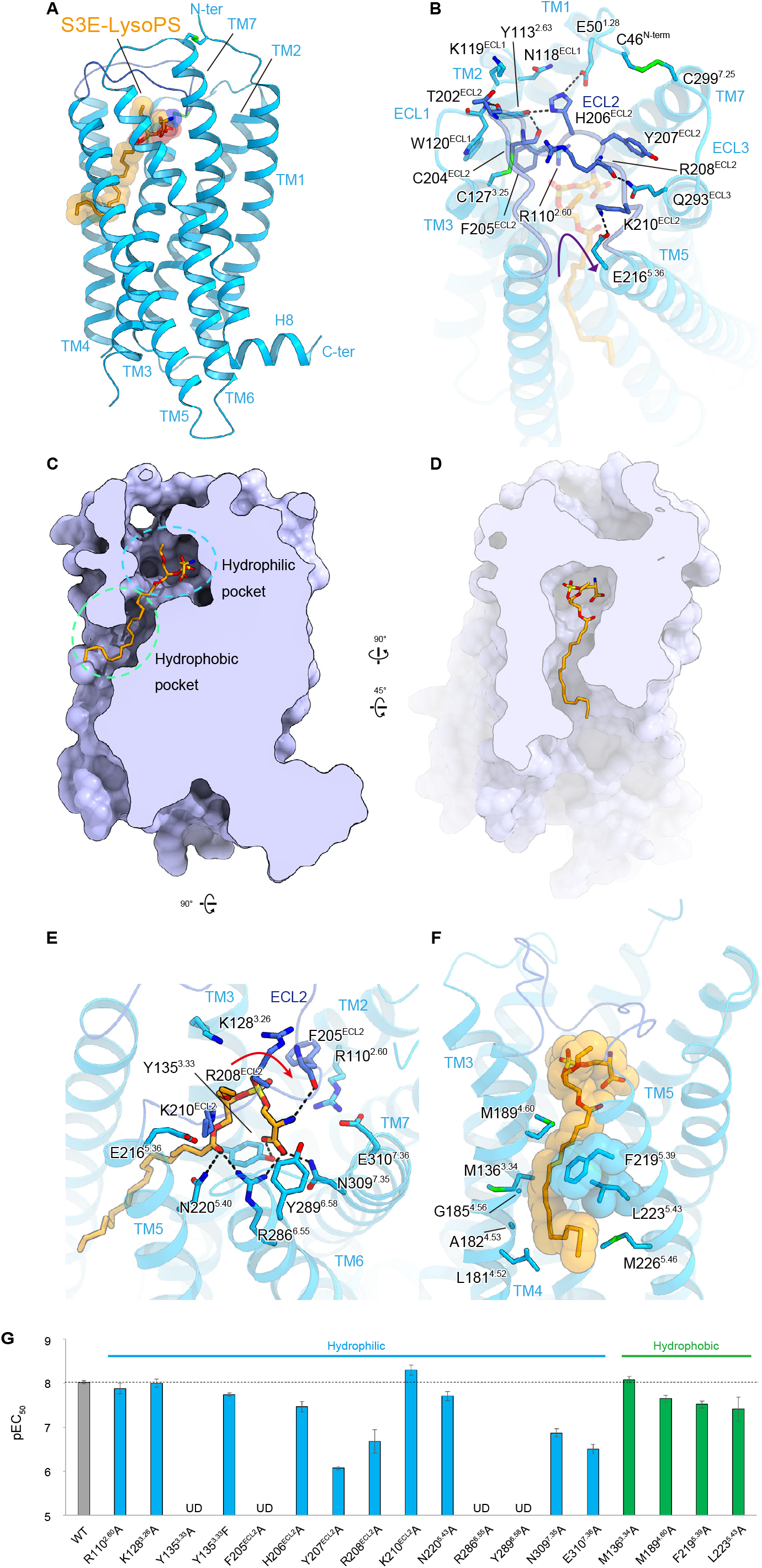
S3E-LysoPS binding mode. (**a**) Overall structure of the S3E-LysoPS-bound receptor. Disulfide bonds are shown as sticks. (**b**) Interactions between extracellular loop (ECL)2 and transmembrane helices (TMs). Black dashed lines indicate hydrogen-bonding interactions. (**c and d**) Cross-sectional views of the ligand-binding pocket, viewed from the membrane plane **(c)** and the extracellular side **(d)**. (**e and f**) Binding mode of S3E-LysoPS in the hydrophilic **(e)** and hydrophobic **(f)** pockets of the receptor.

### The S3E-LysoPS binding mode

The ligand-binding pocket of GPR34 extends from the centre of ECL2 to the middle of TM4–5, forming an approximately 25-Å cleft that is laterally open toward the membrane (Fig. 3c). S3E-LysoPS fits into this cleft, oblique to the receptor (Fig. 3c,d). The ligand-binding pocket further consists of both hydrophilic and hydrophobic pockets. The hydrophilic pocket is the canonical GPCR ligand-binding site, composed of TM2, TM3, TM5–7, and ECL2, whereas the hydrophobic pocket consists of TM4 and TM5. The head group and the acyl chain of S3E-LysoPS fit within the hydrophilic and hydrophobic pockets, respectively.

Y135^3.33^, F205^ECL2^, and Y289^6.58^ create the bottom and sides of the hydrophilic pocket (Fig. 3e), and the head group of S3E-LysoPS fits into the pocket, with a U-shaped conformation. The phosphate group engages in electrostatic interactions with the positively charged residues, R110^2.60^, R208^ECL2^, and K210^ECL2^, but does not form direct interactions, such as hydrogen bonds. In addition, the carboxylate of the serine moiety forms a direct salt bridge with R286^6.55^ and hydrogen bonds with Y135^3.33^ and N309^7.35^. The amine group forms an electrostatic interaction with E310^7.36^ and a hydrogen bond with the backbone carbonyl group of F205^ECL2^. Of note, the oxygen atom in the *sn*-3 position forms a hydrogen bond with N220^5.40^ and R286^6.55^, whereas the ethoxy group in the *sn*-1 position has less contact with the receptor than the other moieties. Overall, these data suggest that GPR34 more firmly recognizes to the amine and carboxylate groups of the serine moiety than the phosphate group.

The hydrophobic pocket consists of a gap between the extracellular halves of TM4 and TM5 (TM4–5 gap; Fig. 3f). This gap is wider than those in the EDG family members of lipid receptors and the phylogenetically related P2Y receptor (P2Y12), owing to different positions of TM4^23–27^ (Extended Data Fig. 3a–g). However, a similarly wide gap is observed in the structure of the non-EDG LPA receptor LPA6^21^ (Extended Data Fig. 3b,c). In GPR34, the gap is composed of hydrophobic residues in TM4 and TM5. Notably, TM5 contains the bulky residues F219^5.39^ and L223^5.43^, whereas the opposite position in TM4 has the small residues A182^4.53^ and G185^4.56^ (Fig. 3f), resulting in formation of an L-shaped hydrophobic pocket (Fig. 3d). The acyl chain of S3E-LysoPS is bent at the cis-9 double bond and fits along the L-shaped pocket. Consequently, the C1–C9 chain is exposed to the membrane environment, consistent with a previous study reporting that GPR34 is activated by LysoPS analogues attached to alkoxy amine chains with various hydrophobic tail lengths^28^. To validate the observed agonist interactions, we mutated receptor residues involved in S3E-LysoPS binding. Within the hydrophilic pocket, alanine mutants of the four aromatic residues Y135^3.33^, F205^ECL2^, Y207^ECL2^, and Y289^6.58^, which are critical for hydrophilic pocket formation, reduced potency of S3E-LysoPS (pEC_50_) by over 100-fold (Fig. 3g, Extended Data Fig. 5, and Extended Data Table 4). In addition, R286^6.55^A mutation abolishes agonist response, whereas alanine mutations of R208^ECL2^, N309^7.35^, and E310^7.36^ reduced potency by approximately 10- to 30-fold. These observations are consistent with the fact that residues involved in head group recognition are highly conserved among vertebrates, indicating their functional importance for LysoPS receptors (Extended Data Fig. 4a,b). In contrast, alanine mutations in the hydrophobic pocket only reduced potency by up to 4-fold (Fig. 3g, Extended Data Fig. 5, and Extended Data Table 4). These hydrophobic pocket residues are less highly conserved compared to those in the hydrophilic pocket, suggesting there is no strict spatial requirement to accommodate the acyl chain.

### Ligand access

The ligand-binding pocket of GPR34 is open toward both the membrane and extracellular space (Fig. 3c,d), suggesting the ligand enters the pocket laterally from the membrane or the extracellular medium^28^. Unlike other lysophospholipids, such as LPA and S1P, which are both present in relatively high amounts as carrier-bound forms in extracellular fluids, LysoPS concentration in extracellular fluids is too low to activate receptors^29^. In addition, LysoPS is produced from PS by the extracellular enzyme PS-PLA_1_ in the outer leaflet of the plasma membrane^17^, with no known pathways for production in the extracellular fluid. Interestingly, when added to the medium, recombinant PS-PLA_1_ protein activates GPR34 at the cellular level (Extended Data Fig. 6a). Under these conditions, LysoPS is not present in the medium but, rather, is associated with cells (Extended Data Fig. 6b,c). When albumin, which can extract lysophospholipids from the membrane, is added simultaneously, PS-PLA_1_-induced GPR34 activation is dramatically weakened (Extended Data Fig. 6a,d), indicating that membrane-associated LysoPS, but not albumin-bound LysoPS, is capable of activating GPR34. Further, a PS-PLA1 S166A mutant only weakly activates GPR34 (Extended Data Fig. 6a). These results, together with a ligand pocket open to the membrane, suggest that LysoPS enters the pocket laterally when produced on the outer leaflet of the plasma membrane by PS-PLA1. Moreover, albumin effectively inhibits M1-induced GPR34 activation in a dose-dependent manner (Extended Data Fig. 6e–g), suggesting lateral access of the synthetic GPR34 agonist, in addition to extracellular access. These findings indicate that the open shape of the ligand-binding pocket is suitable for membrane and extracellular access.

### M1 binding mode

We next analysed the GPR34 structure bound to M1, the metabolically stable S3E-LysoPS analogue^18^. The overall structure of the M1-bound receptor superimposes well on the S3E-LysoPS-bound structure, with a root-mean-square deviation (RMSD) of 0.73 Å (Fig. 4a). M1 binds to the hydrophilic and hydrophobic pockets in a pose similar to that of S3E-LysoPS, and the head groups of M1 and S3E-LysoPS form comparable interactions with the hydrophilic pocket (Fig. 4b). Compared with the S3E-LysoPS-bound form, the extracellular portion of TM4 is displaced outwardly by 2.3 Å (Fig. 4a), due to key differences in the hydrophobic pockets.

**Fig. 4:**
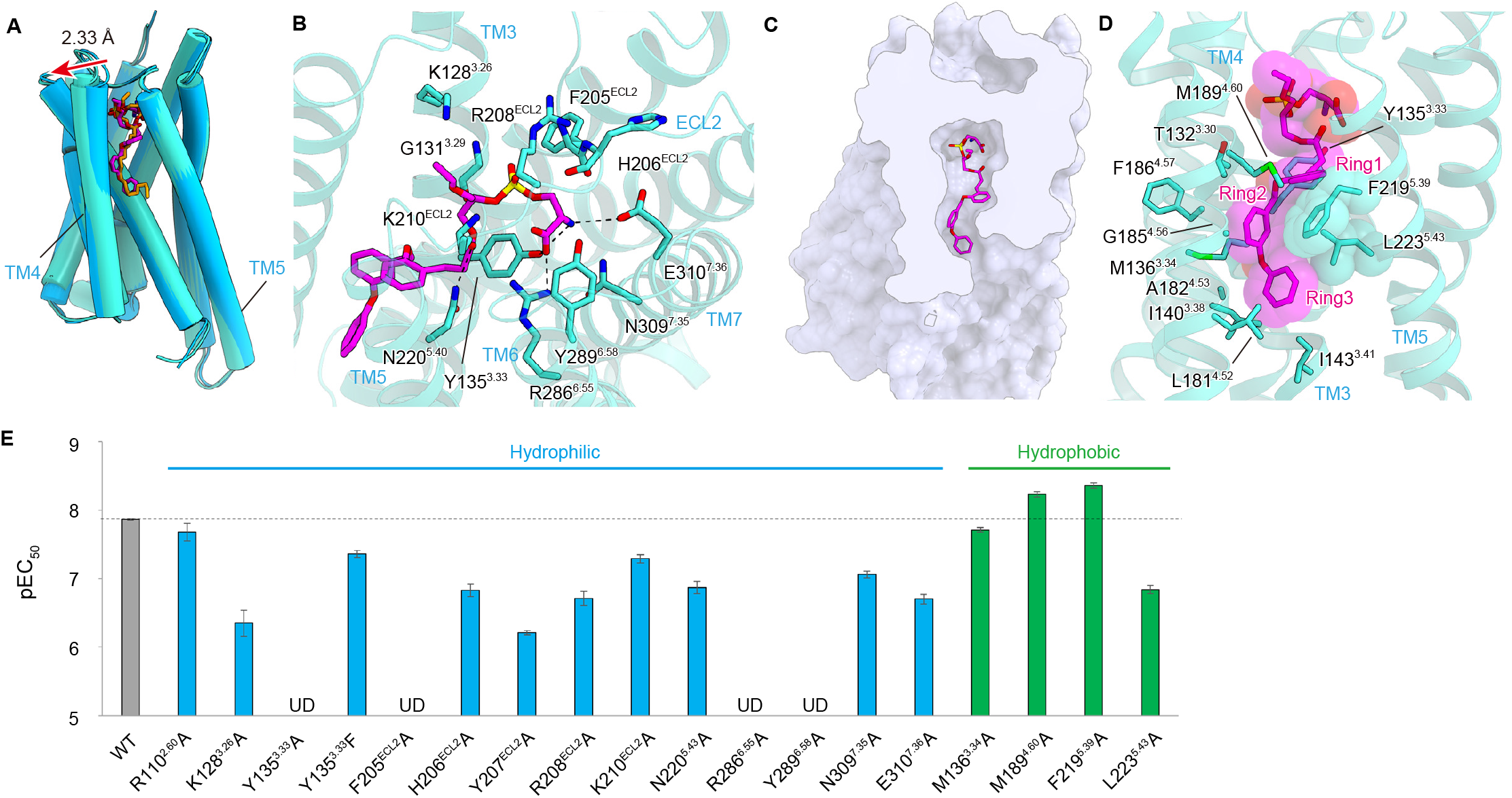
M1 binding mode. (**a**) Superimposition of the GPR34 structures bound to M1 (turquoise) and S3E-LysoPS (blue). **(b**) Binding mode of M1 in the hydrophilic pocket. Black dashed lines indicate hydrogen-bonding interactions. (**c**) Cross-section of the ligand-binding pocket, viewed from the extracellular side. (**d**) Binding mode of M1 in the hydrophobic pocket, highlighting the ring interactions.

Here, the three aromatic rings of M1 (ring1, ring2, and ring3) are accommodated in the TM4–5 gap (Fig. 4c,d), with ring1 and ring3 oriented perpendicular to ring2. The bend between ring2 and ring3 fits along the L-shaped hydrophobic pocket, which superimposes with the position of the Z–C9 double bond of 18:1 in S3E-LysoPS^16^. These aromatic moieties tightly interact with the receptor by stacking interactions with F219^5.39^ and L223^5.43^ (Fig. 4a). Overall, the larger opening of the TM4–5 gap accommodates the bulky aromatic groups of M1 well.

As for S3E-LysoPS, we mutated the residues involved in the M1 binding and observed overall effects is similar to those detected with S3E-lysoPS (Fig. 4e, Extended Data Fig. 5, and Extended Data Table 4). However, within the hydrophobic pocket, F219^5.39^A mutation increased the potency of M1, suggesting its bulkiness is not essential for M1 binding. In contrast, L223^5.43^A mutation reduces potency by 10-fold (Fig. 4e), suggesting the hydrophobic constriction formed by L223^5.43^ provides the necessary environment for M1 binding.

### Validation of agonist binding modes by MD simulations

To validate the observed agonist binding modes, we performed 1-μs MD simulations of receptor–ligand complexes in a 1-palmitoyl-2-oleoyl-sn-glycero-3-phosphocholine (POPC) lipid bilayer environment. During simulations, interactions between phosphoserines and receptors in the cryo-EM structure are stably maintained for both *sn*-3 LysoPS derivatives: S3E-LysoPS and M1 (Fig. 5a,b, Extended Data Fig. 7a, Extended Data Video 1, and Supplementary information). Results further suggest that anionic charge repulsion between the phosphate (PO^−^) and serine CO_2_^-^moieties could contribute to a preference for the U-shaped conformation of the hydrophilic portions of these ligands. This conformation, which is specific to *sn*-3 ligands, may facilitate the stabilizing interaction network between phosphoserine (NH_3_^+^, CO_2_^-^, and PO^−^) charges and the hydrophilic binding-site residues E310^7.36^, N309^7.35^, R286^6.55^, and F205^ECL2^ (Fig. 5a).

**Fig. 5:**
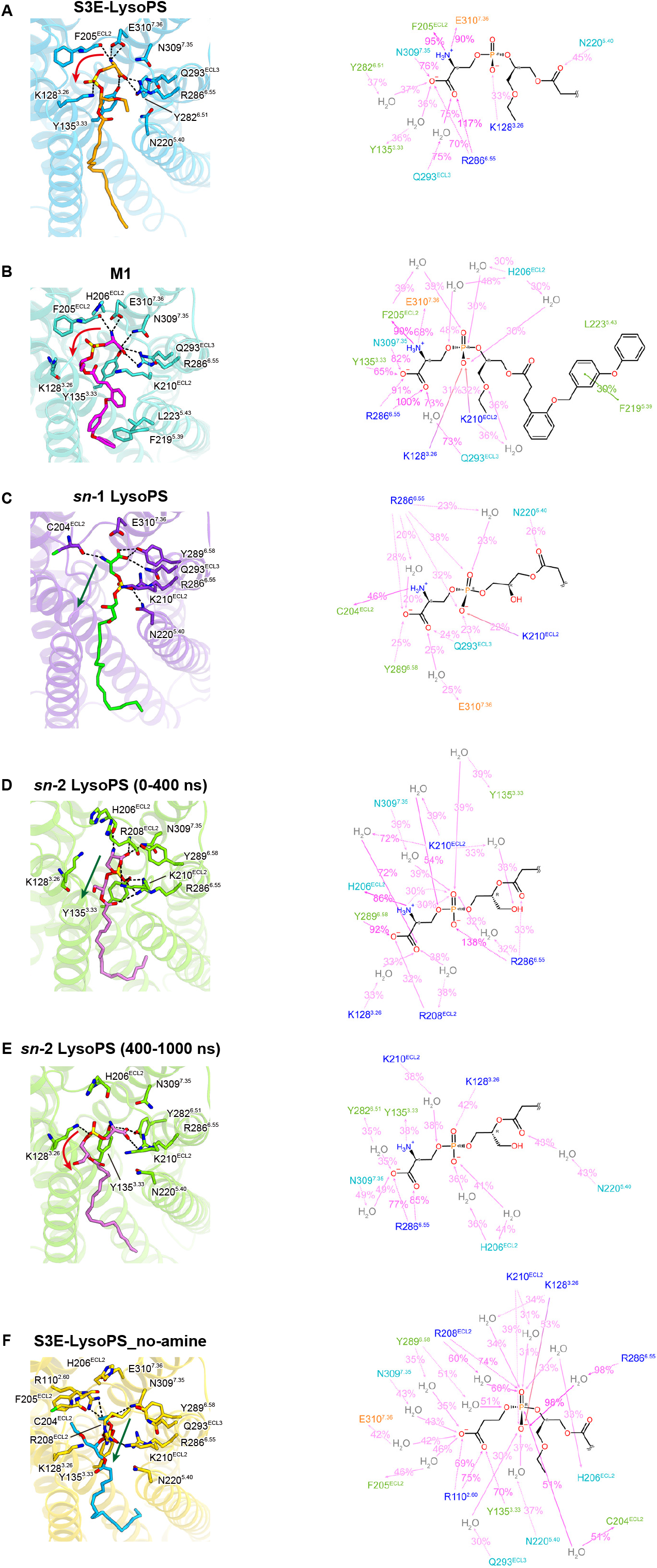
Molecular dynamics (MD) simulations. (**a–f**) Average structures and maintained ligand–protein interactions during 1-μs MD simulations. We illustrate the ligand–protein interactions conserved in over 30% of simulations (Percentage = Number of frames_interaction present_ / Number of frames_total_). Orange, negatively charged residues; blue, positively charged residues; cyan, polar residues; green, hydrophobic residues. The *sn*-1 LysoPS does not form such a stable interaction. Therefore, in this case, we illustrate the interactions conserved in over 20% of simulations. Because the binding pose of *sn*-2 LysoPS is altered significantly after 400 ns, we provide two representations: 0–400 ns and 400–1000 ns.

MD simulations also indicate that M1 forms significant stable interactions with the receptor (Extended Data Fig. 7b,c) and dynamically freezes movement of the extracellular half of TM4 (RMSF_A182-I191_ <0.8 Å) (Extended Data Fig. 7d), whereas S3E-LysoPS allows some movement of this helix (RMSF_A182-I191_ = 1.0–1.8 Å) (Extended Data Fig. 7e). Additionally, distances between residues in TM4 and TM5 are constant in the M1-bound receptor but varied in the S3E-bound receptor. This suggests that the tight hydrophobic interactions of M1 hold TM4 in a more open conformation (Extended Data Fig. 7f–h). Consistently, stacking interactions with F219^5.39^ and L223^5.43^ are stably maintained during MD simulation (Fig. 5b). Because TM4 is separate from, and undergoes few interactions with, other TMs, it has high flexibility, as shown in MD simulation of the S3E-bound receptor (Extended Data Fig. 7e). This flexible nature would allow lateral access of the ligand and induced fit upon M1 binding. To date, TM4 has been somewhat ignored in GPCR studies because it does not face the canonical ligand-binding pocket; however, it plays an essential role in drug recognition by GPR34.

### Regioselectivity of LysoPS species binding to GPR34

We further performed docking simulations with natural *sn*-1 and *sn*-2 LysoPS species and compared dynamics with those of the synthetic *sn*-3 analogue S3E-LysoPS (bioactivities: *sn*-3 > *sn*-2 >> *sn*-1; Fig. 1b,c and Extended Data Table 1 and 2). For *sn-*1 LysoPS (18:1), we found that the phosphoserine occupies a binding position with high probability that is distinct from its position in S3E-LysoPS and M1 (Fig. 5c, Extended Data Fig. 7a, Extended Data Video 1, and Supplementary information). That is, the phosphoserine adopts a straight-line shape rather than the U-shape observed in S3E-LysoPS and M1. Further, in this case, instead of the carboxyl group (as in S3E-LysoPS or M1), the phosphate group forms hydrogen bonds with N220^5.40^ and R286^6.55^ (Fig. 5c). For *sn*-2 LysoPS (18:1), the initial docking pose is similar to that of *sn*-1 LysoPS (Fig. 5d, Extended Data Fig. 7a, and Extended Data Video 1). However, it gradually switches to an S3E-LysoPS-like pose after 400 ns of simulation (Fig. 5e). This conformational mode, which shares a common interaction network with the *sn*-3 ligands (Fig. 5a,b), is maintained throughout the following 600 ns (Fig. 5e, Extended Data Fig. 7a, and Extended Data Video 1). Critically, the above observation suggests that the U-shaped conformation of the hydrophilic head may represent the active form of the ligand and is essential for preserving the hydrophilic interaction network with the protein (E310^7.36^, N309^7.35^, R286^6.55^, and backbone carbonyl of F205^ECL2^). Finally, MD simulations with a ligand lacking the amine group in the serine head (S3E-LysoPS-no_amine) result in a binding mode switch from the *sn*-3 type to an *sn*-1-type-like mode within 300 ns (Fig. 5f, Extended Data Fig. 7a, and Extended Data Video 1), wherein the hydrophilic head adopts a straight-line shape. This is consistent with our previous findings regarding GPR34 ligand specificity; that is, only LysoPS species with a phosphoserine head group can activate this receptor^1^. This likely results from the fact that the hydrophilic binding sites of GPR34 contain an assembly of charged amino acid residues that can efficiently interact with the charged phosphoserine (Fig. 5a).

### Receptor activation and G_i_ coupling

Although the inactive GPR34 structure has not yet been solved, our active GPR34 structure provides mechanistic insight into receptor activation (Fig. 6a). In the homologous receptor P2Y12, positively charged residues, such as R256^6.55^, form salt bridges with the phosphate groups of nucleic acids (Extended Data Fig. 8a), causing the 4-Å inward shift of TM6^23,30^. Likewise, S3E-LysoPS tightly interacts with R286^6.55^ (Extended Data Fig. 8b), which is also observed in MD simulations of the *sn*-1 and *sn*-2 LysoPS-bound forms (Fig. 5c–e). Below R286^6.55^, Y282^6.51^ hydrogen bonds with Y135^3.33^, and H283^6.52^ forms a π-stacking interaction with F227^5.47^ (Fig. 6b). Analogous to P2Y12, interaction between LysoPS and R286^6.55^ could induce an inward displacement of the extracellular portion of TM6, leading to formation of the central core interaction.

**Fig. 6:**
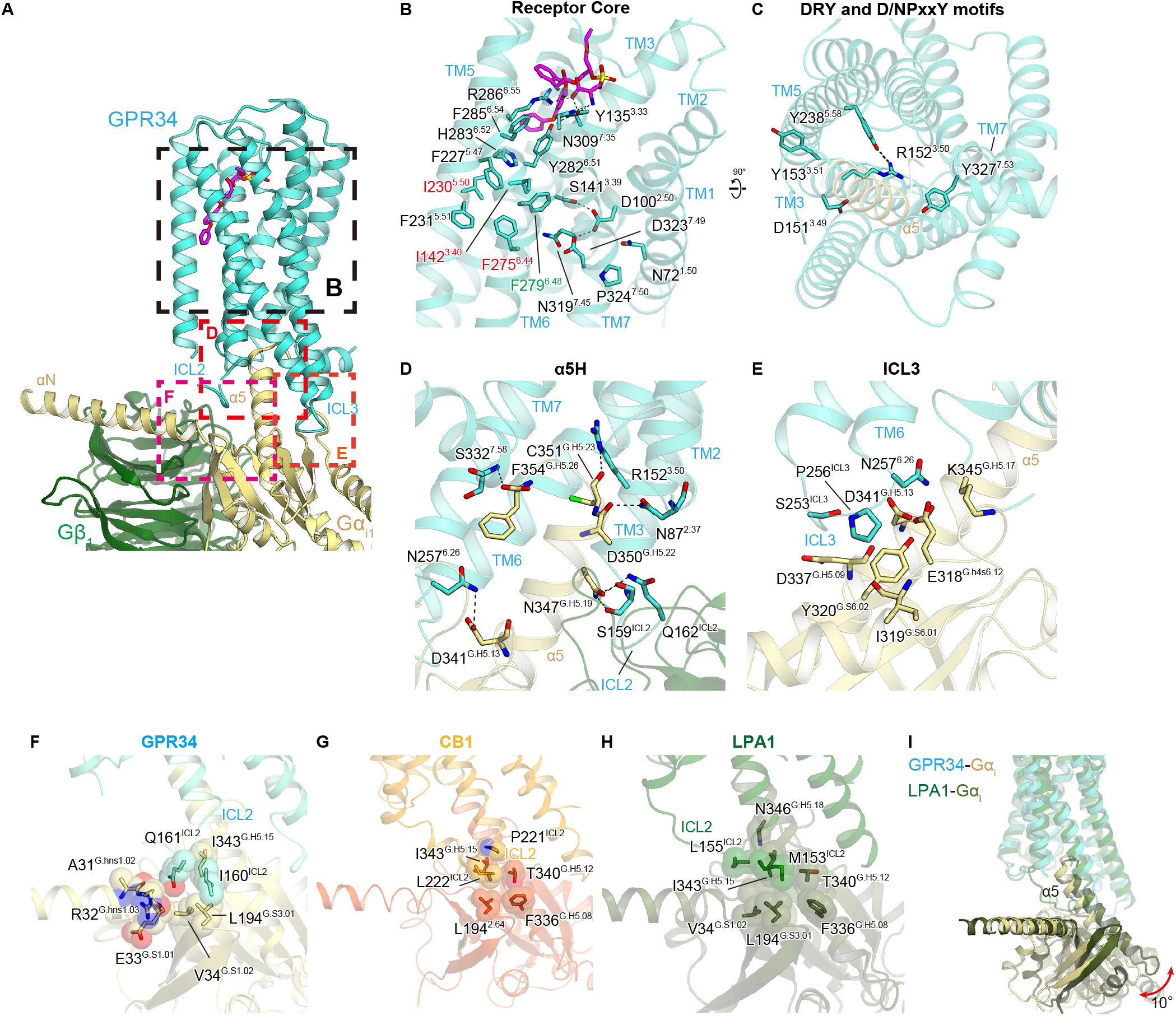
G-protein activation. (**a**) G-protein interface. (**b**) Interactions at the receptor core. (**c**) DRY and N/DPxxY motifs. Black dashed lines indicate hydrogen bonds. (**d**) Main hydrogen-bonding interactions between the receptor and the α5-helix of Gαi1. (**e**) Interactions between intracellular loop (ICL)3 and G_i_. (**f–h**) Interactions between ICL2 and G_i_ in GPR34 **(f)**, CB1 (PDB 6N4B) **(g)**, and lysophosphatidic acid (LPA)1 (PDB 6N4B) **(h)**. (**i**) Comparison of the G-protein positions in the GPR34 and LPA1 (PDB 7YU4) structures.

In most class A GPCRs, ligand binding rearranges hydrophobic contacts in the conserved P-I-F and CWxP motifs in the centres of TMs 3-5-6, leading to receptor activation^22,31^. P^5.50^ is not conserved, as described above, and W^6.48^ is replaced by F279^6.48^ (Fig. 6b). However, these hydrophobic residues are tightly packed together with the nearby phenylalanine (Fig. 6b). A polar interaction network exists in the middle parts of TMs 1, 2, and 7, including the conserved D323^7.49^ in the N/DPxxY motif and D100^2.50^ (Fig. 6b). Formation of these interactions upon ligand binding creates an open cavity for TM5–6 on the intracellular side. Within this intracellular cavity, R152^3.50^ and Y327^7.53^ in the conserved DRY and N/DpxxY motifs are directed toward the centre of the transmembrane bundle (Fig. 6c), facilitating interactions with the C-terminal residues in the α5-helix of G_i_.

The cavity closely contacts the C-terminal α5-helix, and the cytoplasmic loops, particularly ICL2 and ICL3, may contribute to G protein interactions. Specifically, the α5-helix C-terminus forms a hydrogen bond with the backbone amide of S332^7.58^ (Fig. 6d). Moreover, R152^3.50^ hydrogen bonds with the backbone carbonyl of C351^G.H5.23^ (superscript indicates the common Gα numbering [CGN] system), as typically observed in other GPCR-Gi complexes^32^. The short ICL3 forms van der Waals interactions with the α5-helix and β-sheet of G_i_, and N257^6.26^ hydrogen bonds with E318^G.H4S6.12^ and D341^G.H5.13^ (Fig. 6e). The most characteristic feature of the GPR34-G-protein interface is the interaction at ICL2 (Fig. 6f), which is located at the root of the α5-helix, and thus has a significant effect on its orientation. In most receptor-Gs and -G_i_ complex structures^33–36,32,37^, bulky hydrophobic residues in the α-helix of ICL2 fit into hydrophobic pockets formed by L194^G.S3.01^, F336^G.H5.08^, T340^G.H5.12^, and I343^G.H5.15^ in the Gαi subunits (Fig. 6g). In some cases (*i.e*., LPA1-G_i_ complex)^26,25^, ICL2 adopts a disordered conformation, but M153^ICL2^ still fits into the hydrophobic pocket of G_i_ (Fig. 6h). Conversely, in GPR34, the hydrophobic residues in ICL2 do not fit in the pocket (Fig. 6f). Rather, I160^ICL2^ and Q161^ICL2^ form superficial interactions with the αN, α5-helix, and β-sheet of G_i_. Due to these differences, the α5-helix in the GPR34-G_i_ complex is 10° perpendicular to the receptor, compared to its position in the LPA1-G_i_ complex (Fig. 6i). Overall, the GPR34-G_i_ coupling interaction is unique relative to that of other GPCR-G_i_ complexes and extends the diverse binding modes observed for G_i_ compared to G_s_.

### Insight into superagonism of M1

Structures of the essential motifs and G-protein coupling interactions are similar in the M1- and S3E-LysoPS-bound receptors (Extended Data Fig. 8c). However, because M1 functions as a superagonist, unlike S3E-LysoPS, it forms significant stable interactions with the receptor (Extended Data Fig. 7b,c), which can freeze motion of the TM4–5 gap (Extended Data Fig. 7d,e). Moreover, in 1-μs MD simulations, RMSF values for the intracellular sides of TM5–7 and H8 are larger in the M1-bound GPR34 than the S3E-LysoPS-bound structure. It is possible that M1 prefers the active state during MD simulation, where TM6 and TM7 move farther from the centre of the TM bundle (Extended Data Fig. 8d), thus enabling higher activity for G-protein activation.

## Discussion

In summary, the S3E-LysoPS-bound cryo-EM structure revealed acyl chain accommodation in the TM4–5 gap (Fig. 3f), a unique characteristic among lipid-sensing GPCRs. MD simulations based on the structure further showed that head groups of *sn*-2 and *sn*-3 LysoPS can adopt U-shapes and form tight interactions with charged groups in the hydrophilic pocket (Fig. 5a,e). Notably, the amine and carboxylate of the serine moiety are tightly recognized by E310^7.36^ and R286^6.55^ (Fig. 5a), respectively, accounting for serine-specific recognition by the LysoPS receptor GPR34. In contrast, the head group of *sn*-1 LysoPS can only adopt a straight shape and does not form such stable interactions (Fig. 5c), consistent with the limited GPR34 activity of *sn*-1 LysoPS species. Together with the results of our cell-based assay (Fig. 1b) and previous reports^1^, this structural study therefore shows that *sn*-2 LysoPS is the physiological ligand of GPR34. However, due to the short half-life of *sn*-2 LysoPS (easy transition to *sn*-1) and the necessity of enzymatic production, its preparation might be complicated, resulting in low purity. This is one possible reason for the controversy regarding whether LysoPS can activate GPR34. Moreover, the *sn*-2 acyl selectivity may control the signal duration time of GPR34, which might be shorter than those of other lipid receptors. Critically, the metabolically stable agonist M1 mimics the acyl chain conformation of LysoPS in the TM4–5 gap, forming quite stable interactions with the receptor, and thus can function as a superagonist. Overall, we anticipate that detailed SAR information and physiological functional data in this and future studies will enable therapeutic targeting of GPR34.

The binding mode of the acyl chain in GPR34 differs substantially from that in other lysophospholipid receptors (Extended Data Fig. 3h–m). For example, EDG receptors accommodate the acyl chain within a transmembrane pocket (Extended Data Fig. 3i–l), whereas GPR34 does so in the TM4–5 gap. However, the crystal structure of the non-EDG LPA receptor LPA6 suggests is may also accommodate the acyl chain in the TM4–5 gap (Extended Data Fig. 3c). Because GPR34 and the non-EDG family LPA receptors are homologous to P2Y receptors, this suggests that acyl chain accommodation in the TM4–5 gap is a common feature of P2Y-like lipid receptors. It is unclear, however, why EDG and P2Y-like lipid receptors accommodate acyl chains differently. P2Y receptors are a family of purinergic G protein-coupled receptors activated by nucleotides^3^, such as adenosine triphosphate. P2Y-like lipid receptors evolved from P2Y family members to receive the acyl chain linked to the phosphate head. Notably, compared to EDG receptors, the phosphate-binding site of P2Y receptors is buried inside the transmembrane bundle (Extended Data Fig. 3M). We therefore propose that due to limited space, P2Y receptors used the membrane-facing hydrophobic region to evolve as lipid receptors.

The LysoPS receptors P2Y10 and GPR174 share 50% sequence identity, whereas GPR34 is more distantly related, and there is no obvious conservation of residues required for LysoPS binding (Extended Data Fig. 9a). In addition, ECL2, which is essential for ligand recognition, is predicted to differ at the secondary structure level in GPR34 relative to P2Y10 and GPR174. To better understand the relationships between these receptors, we can leverage recent AlphaFold-2 (AF2)^38^-predicted structures to perform a structural comparison of the three LysoPS receptors. Critically, both the cryo-EM and predicted structures of GPR34 contain identical TM4–5 gaps (Extended Data Fig. 9b–d), although the extracellular and intracellular faces are different, suggesting it is possible to compare TM4–5 gaps in the AF2-predicted structures. In doing so, we find that the TM4–5 gap is similar in all three LysoPS receptors (Extended Data Fig. 9b–f), implying that in all cases, it accommodates the acyl chain. However, F219^5.39^ and L223^5.43^, which form L-shaped hydrophobic pockets in GPR34, are not conserved in P2Y10 and GPR174, and thus, the TM4–5 gaps in these receptors adopt more open conformations (Extended Data Fig. 9e,f). Previous structure–activity relationship studies have further shown that acyl chain modifications alter the ligand selectivity for LysoPS receptors. As observed for the aromatic rings of M1, recognition of the L-shaped pocket is critical for GPR34 selectivity.

## Acknowledgments

We thank K. Ogomori and C. Harada for technical assistance. This work was supported by grants from the JSPS KAKENHI, grant numbers 21H05037 (O.N.), 22K19371 and 22H02751 (W.S.), and 22H00438 (J.A.); the ONO Medical Research Foundation (W.S.); The Kao Foundation for Arts and Sciences (W.S.); The Takeda Science Foundation (W.S.); The Uehara Memorial Foundation (W.S.); the Kobayashi Foundation, Osaka, Japan (T.O.); the KOSÉ Cosmetology Research Foundation (T.O.); the Japan Agency for Medical Research and Development (AMED), grant numbers 22ck0106533h0003 (J.A., O.N. and T.O.) and 21gm0010004h9905 (J.A.); and the Platform Project for Supporting Drug Discovery and Life Science Research (Basis for Supporting Innovative Drug Discovery and Life Science Research (BINDS)) from AMED, grant numbers JP19am01011115 (support no. 1109, O.N.). L.C. wishes to also thank the Otsuka Toshimi Scholarship Foundation, Osaka, Japan.

## Author contributions

T.I. expressed, purified, and prepared grids of the S3E-LysoPS bound GPR34-G_i_ complex, with assistance from K.K. and H.O. Y.K. performed the cryo-EM analysis of the S3E-LysoPS-bound complex. T.I. and W.S. performed single-particle analysis and model building of the S3E-LysoPS-bound complex. R. K. performed the structural study of the M1-bound receptor, with assistance from T.T., F. K. S., and W.S. W.S. initially screened and established the protocol for sample preparation. A.U., S.Y. and J.O. performed the TGFα shedding assay, the cAMP assay, and LC-MS/MS analysis, and wrote part of the manuscript. H.K. assisted with lipid preparation for the assay. J.A. supervised most of the biological experiments and wrote the biological part of the manuscript. L.C. and T.O. performed and oversaw the MD simulations and wrote the simulation part of the manuscript. The manuscript was mainly prepared by W.S., R.K., T.I., T.O., and J. A., with input from all authors and assistance from O.N.

## Data and materials availability

Density maps and structure coordinates have been deposited in the Electron Microscopy Data Bank (EMDB) and the PDB, with accession codes EMD-34512 and PDB 8H6W for the S3E-LysoPS-GPR34-G_i_ complex; EMD-34513 and PDB 8H6X for the S3E-LysoPS-GPR34-G_i_ complex (Receptor focused); EMD-34514 and PDB 8H6Y for the M1-GPR34-G_i_ complex; EMD-34515 and PDB 8H6Z for the M1-GPR34-G_i_ complex (Receptor focused).

## Methods

### Preparation of *sn* −1 LysoPS (18:1) and *sn*-2 LysoPS (18:1)

The *sn*-1 LysoPS (18:1) and *sn*-2 LysoPS (18:1) agonists were prepared as described previously^39^. Briefly, di-oleoyl (18:1) phosphatidylserine (PS) (di-18:1-PS) from Avanti Polar Lipids (Alabaster, AL, USA) were digested with *Rhizomucor miehei* lipase, which is known to have intrinsic phospholipase A1 (PLA1) activity. The resulting *sn*-2 LysoPS (18:1) was stabilized by bringing the solvent to a mild acidic pH of 4.0 to prevent the acyl migration reaction. The PLA1 reaction mixture containing *sn*-2 LysoPS (18:1), di-18:1-PS, and oleic acid was then subjected to C18-based reverse-phase cartridge column chromatography to separate *sn*-2 LysoPS (18:1). After obtaining a pure *sn*-2 LysoPS (18:1) preparation, the solvent was changed to alkaline conditions (pH 9.0) to facilitate the acyl migration reaction. After neutralisation, the resulting LysoPS was used as *sn*-1 LysoPS (18:1). We assessed the purities of *sn*-2 LysoPS (18:1) and *sn*-1 LysoPS (18:1) preparations by liquid chromatography–tandem mass spectrometry (LC-MS/MS) and confirmed that these were >90% pure.

### Transforming growth factor (TGF)-α shedding assay

The TGF-α shedding assay was performed as described previously^40^. Briefly, HEK293A cells were seeded in 6-well plates at a density of 4×10^5^ cells/well and cultured for 1 day in a 5% CO_2_ incubator at 37°C. Cells were then transfected with a mixture of plasmids encoding alkaline phosphatase-tagged (AP)-TGFα (500 ng), human GPR34 (200 ng), and Gαq/i1, a chimeric Gα protein (100 ng), using polyethyleneimine (Polysciences, Inc., Warrington, PA, USA), and cultured for an additional day. Negative control cells were transfected with empty plasmid instead of the GPR34-encoding plasmid. Transfected HEK293A cells were harvested with 0.05% trypsin/EDTA and seeded in 96-well plates (2.5×10^4^ cells/well). Cells were then treated with various LysoPS agonists and a PS-PLA_1_ recombinant protein in the presence of Ki16425, a lysophosphatidic acid (LPA)_1/3_ antagonist (final concentration, 3 mM), for 60 min at 37°C. After centrifugation, the supernatant was transferred to another plate and 10 mM *p*-NPP was added to both the supernatant and cell plates, at a volume of 80 mL/well. Finally, the optical density at 405 nm (OD_405_) was measured with a SpectraMAX ABS Plus (Molecular Devices, San Jose, CA, USA), before and after incubation at room temperature. AP-TGFα release was calculated as follows:

AP-TGFα release (%) = DOD405_Sup_ / (DOD405_Sup_ + DOD405_Cell_) × 100 × 1.25. In this equation, we multiply by 1.25 to convert the amount of AP-TGFα in the transferred supernatant (80 mL) to the amount of AP-TGFα in total supernatant (100 mL). We then calculated GPCR activation as: GPCR activation (%) = AP-TGFα release under stimulated conditions (%) – AP-TGFα release under unstimulated conditions (%).

GPCR activation levels were fit to four-parameter sigmoidal concentration–response curves, using Prism9 software (GraphPad, San Diego, CA, USA), and pEC_50_ and E_max_ values were obtained from the curves.

### cAMP assay

The cAMP assay was performed with GloSensor cAMP Biosensor (Promega, Madison, WI, USA), as previously described^40^. Briefly, HEK293A cells were seeded and cultured as described for the TGF-α shedding assay above. Cells were then transfected with a mixture of plasmids encoding GloSensor-22F (1 mg) and human GPR34 (200 ng), using polyethyleneimine (Polysciences), and cultured for an additional day. Negative control cells were transfected with empty plasmid instead of the GPR34-coding plasmid. Transfected HEK293A cells were harvested in Dulbecco’s Phosphate-Buffered Saline (D-PBS), containing 2 mM EDTA, and resuspended in 0.01% bovine serum albumin (BSA)/HBSS. Cells were then seeded in half-area white 96-well plates (3.5×10^4^ cells/well) and loaded with D-Luciferin (final concentration, 2 mM). After incubation in the dark for 2 h at room temperature, basal luminescence was measured by a SpectraMAX L microplate reader (Molecular Devices). Cells were then treated with forskolin (final concentration, 10 mM) and various LysoPS in the presence of Ki16425, an LPA_1/3_ antagonist (final concentration, 3 mM), and post-stimulus luminescence was kinetically measured for 20 min at room temperature. We then calculated cAMP (% Forskolin stimulation) as follows: post-stimulus luminescence was normalized by dividing the raw values by the basal luminescence, and normalized luminescence in both agonist and forskolin-treated conditions was divided by that in forskolin-only treated conditions. To obtain pEC_50_ and E_max_ values, cAMP signals were fitted to four-parameter sigmoidal concentration–response curves, using Prism9 software (GraphPad).

### GPR34 activation by PS-PLA_1_

Recombinant PS-PLA1 was prepared as described previously^41^, with minor modifications. In brief, HEK293A cells were transfected with plasmid (1 mg) encoding wild-type (WT) or S166A-mutant mouse PS-PLA_1_, using Lipofectamine 2000 (Thermo Fisher Scientific, Waltham, MA, USA). Negative control cells were transfected with the empty plasmid. After 4 h, the medium was changed to Opti-MEM, and the HEK293A cells were cultured for 72 h in a 5% CO_2_ incubator at 37°C. The culture supernatant was then collected and centrifuged at 400 ×*g* for 5 min, and the resulting supernatant was used as recombinant PS-PLA_1_. For evaluation of GPR34 activation by PS-PLA_1_, the TGFα shedding assay was performed, as described above, using recombinant PS-PLA_1_ protein in place of LysoPS.

### Sample preparation for LC-MS/MS analysis

The amount of LysoPS in HEK293A cells and in supernatant from HEK293A cells stimulated by recombinant PS-PLA_1_ was determined by LC-MS/MS analysis. Samples for LC-MS/MS analysis were prepared as described previously^42^. Briefly, HEK293A cells were stimulated by PS-PLA_1_ as described above, and the entire supernatant was collected. Cells were treated with ice-cold acidic MeOH, containing 100 nM 17:0-LPA, and incubated for 10 min at room temperature; LysoPS dissolved in MeOH was then collected. For supernatant samples, 10 mL of the collected supernatant was added to 90 mL of acidic MeOH, containing 111 nM 17:0-LPA. Both cell and supernatant samples were passed through a filter with a 0.2 mM pore size and a 4 mm inner diameter and subjected to LC-MS/MS analysis, as described below.

### LC-MS/MS analysis

LC-MS/MS analysis was performed as described previously^42^, using an LC-MS/MS system consisting of a Vanquish HPLC system and a TSQ Altis™ Triple-Stage Quadrupole Mass Spectrometer (Thermo Fisher Scientific). For HPLC, samples were separated in the L-column2 (100 mm×2 mm, 3-mm particle size, CERI), using a gradient solution consisting of solvent A (5 mM ammonium formate in water, pH 4.0) and solvent B (5 mM ammonium formate in acetonitrile, pH 4.0) at 200 ml/min. LysoPS was then monitored in the negative ion mode, using MS/MS. At MS1, the m/z values of [M+H]+ ion for LysoPS were selected. At MS3, lysophosphatidic acid fragments derived from LysoPS were detected. The amount of LysoPS in samples was calculated based on the standard curve of 18:1-LysoPS.

### Preparation of anti-GPR34 serum

Anti-GPR34 serum was obtained by performing DNA immunization^43^ of *Gpr34*-knock-out (KO) mice, to ensure that GPR34 was recognized as an antigen. Mice were intramuscularly injected with pCAGGS (100 mg) plasmid, which encodes a mouse GPR34–GroEL fusion protein, and electroporated *in vivo*. Immunization was performed five times in total, once every two weeks. Two days after the last immunization, mice were boosted by intrasplenic administration of mouse GPR34-expressing HEK293T cells (2×10^7^ cells), and serum was collected 3 days later. This serum was used as anti-GPR34 serum.

### Evaluation of GPR34 mutant expression

Expression of GPR34 mutants was measured by flow cytometry. In brief, HEK293A cells were transfected with human GPR34-encoding plasmid (250 ng), using polyethyleneimine. Cells were then suspended in 200 ml of D-PBS, containing 2 mM EDTA, and dispensed into 96-well V-bottom plates. After centrifugation for 1 min at 700 ×*g*, the cells were suspended in 200 ml/well of FACS buffer (D-PBS, containing 0.5% BSA and 2 mM EDTA) and incubated for 30 min on ice. Cells were then centrifuged again for 1 min at 700 ×*g*, resuspended in 25 ml/well of anti-human GPR34 serum (1/100 diluted), and incubated for 30 min on ice. After centrifugation for 1 min at 700 ×*g*, cells were washed with D-PBS, resuspended in 25 ml/well of goat anti-mouse IgG conjugated with Alexa488 (Thermo Fisher Scientific, 10 mg/ml), and incubated for 15 min on ice. Cells were then centrifuged a final time for 1 min at 700 ×*g*, washed with D-PBS, and resuspended in 150 ml/well of D-PBS, containing 2 mM EDTA. Flow cytometry analysis was performed with the BD FACSLyric Flow Cytometry System (BD Biosciences, Franklin Lakes, NJ, USA), and the data were analysed by FlowJo Software (FlowJo, Ashland, OR, USA).

### Expression and purification of human GPR34

GPR34 was subcloned into a modified pEG Bacmam vector^44^, with an N-terminal haemagglutinin signal peptide, followed by the Flag-tag epitope (DYKDDDD), and a C-terminal tobacco etch virus (TEV) protease recognition site, followed by an enhanced green-fluorescent protein (EGFP)-His tag ^45^. Recombinant baculovirus was prepared using the Bac-to-Bac baculovirus expression system and *Spodoptera frugiperda* Sf9 insect cells (Thermo Fischer Scientific). The receptor was expressed in HEK293S GnTI-(N-acetylglucosaminyl-transferase I-negative) cells, obtained from the American Type Culture Collection (ATCC; catalogue no. CRL-3022).

To purify the S3E-LysoPS-bound receptor, harvested cells were solubilized in buffer, containing 20 mM Tris-HCl, pH 8.0, 150 mM NaCl, 1% lauryl maltose neopentyl glycol (LMNG; Anatrace, Maumee, OH, USA), 0.1% cholesteryl hemisuccinate (CHS), 10% glycerol, and 10 μM S3E-LysoPS, for 1 h at 4°C. The supernatant was separated from the insoluble material by ultracentrifugation at 180,000 ×*g* for 30 min and then incubated with Flag-M1 resin (Merck & Co., Inc., Rahway, NJ, USA,) for 2 h. Bound resin was washed with 20 column volumes of buffer, containing 20 mM Tris-HCl, pH 8.0, 500 mM NaCl, 0.05% glyco-diosgenin (GDN; Anatrace), 1 μM S3E-LysoPS, 10% glycerol, and 5 mM CaCl_2_. The receptor was then eluted in 20 mM Tris-HCl, pH 8.0, 150 mM NaCl, 0.1% GDN, 1 μM S3E-LysoPS, 10% glycerol, 5 mM EDTA, and 0.15 mg ml^−1^ Flag peptide. The receptor was concentrated and loaded onto a Superdex 200 10/300 column in 20 mM Tris-HCl, pH 8.0, 150 mM NaCl, 0.01% GDN, and 1 μM agonist, and peak fractions were pooled and frozen in liquid nitrogen.

To purify the M1-bound receptor, harvested cells were disrupted by sonication in buffer containing 20 mM Tris-HCl, pH 8.0, 200 mM NaCl, and 10% glycerol. The crude membrane fraction was collected by ultracentrifugation at 180,000 ×*g* for 1 h, and the membrane fraction was solubilized in buffer, containing 20 mM Tris-HCl, pH 8.0, 200 mM NaCl, 2% LMNG (Anatrace), 0.4% CHS, and 10 μM M1, for 1 h at 4°C. The supernatant was separated from the insoluble material by ultracentrifugation at 180,000 ×*g* for 20 min and incubated with TALON resin (Clontech, Mountain View, CA, USA) for 30 min. Bound resin was washed with ten column volumes of buffer, containing 20 mM Tris-HCl, pH 8.0, 500 mM NaCl, 0.05% GDN, 1 μM M1, and 15 mM imidazole. The receptor was then eluted in buffer, containing 20 mM Tris-HCl, pH 8.0, 500 mM NaCl, 0.05% GDN, 1 μM M1, and 200 mM imidazole. The receptor was concentrated and loaded onto a Superdex 200 10/300 Increase size-exclusion column, equilibrated in buffer containing 20 mM Tris-HCl, pH 8.0, 150 mM NaCl, 0.01% GDN, and 1 μM M1. Peak fractions were pooled and frozen in liquid nitrogen.

### Expression and purification of the G_i_ heterotrimer

The G_i_ heterotrimer was expressed and purified using the Bac-to-Bac baculovirus expression system, according to the method reported previously^32^. In brief, Sf9 insect cells were infected at a density of 3–4×10^6^ cells ml^−1^ with a 100th volume of two viruses, one encoding the WT human Gαi1 subunit and the other encoding the WT bovine Gγ_2_ subunit and the WT rat Gβ_1_ subunit containing a His8 tag followed by an N-terminal tobacco etch virus (TEV) protease cleavage site. Infected Sf9 cells were incubated in Sf900II medium at 27°C for 48 h and collected by centrifugation at 6,200 ×*g* for 10 min. The collected cells were then lysed in buffer containing 20 mM Tris, pH 8.0, 150 mM NaCl, and 10% glycerol. The Gαi1β1γ2 heterotrimer was solubilized at 4°C for 1 h in buffer containing 20 mM Tris, pH 8.0, 150 mM NaCl, 10% glycerol, 1% (w/v) *n*-dodecyl-beta-D-maltopyranoside (DDM; Anatrace), 50 μM GDP (Roche), and 10 mM imidazole. The soluble fraction containing Gi1 heterotrimers was then isolated by ultracentrifugation at 186,000 ×*g* for 20 min, and the supernatant was mixed with Ni-NTA Superflow resin (QIAGEN, Hilden, Germany) and stirred at 4°C for 1 h. Bound resin was washed with 10 column volumes of buffer, containing 20 mM Tris pH 8.0, 150 mM NaCl, 0.02% DDM, 10% glycerol, 10 μM GDP, and 30 mM imidazole. Gi1 heterotrimers were then eluted with two column volumes of buffer, containing 20 mM Tris, pH 8.0, 150 mM NaCl, 0.02% (w/v) DDM, 10% (v/v) glycerol, 10 μM GDP and 300 mM imidazole. The eluted fraction was dialysed overnight at 4°C against 20 mM Tris, pH 8.0, 50 mM NaCl, 0.02% DDM, 10% glycerol, and 10 μM GDP. To cleave the histidine tag, TEV protease was added during the dialysis. The dialysed fraction was then incubated again with Ni-NTA Superflow resin at 4°C for 1 h. The flow-through was collected and purified by ion-exchange chromatography on a HiTrap Q HP column (GE Healthcare Life Sciences, Chicago, IL, USA), using Buffer I1 (20 mM Tris, pH 8.0, 50 mM NaCl, 0.02% DDM, 10% glycerol, and 1 μM GDP) and Buffer I2 (20 mM Tris, pH 8.0, 1 M NaCl, 0.02% DDM, 10% glycerol, and 1 μM GDP).

### Expression and purification of scFv16

The gene encoding scFv16 was synthesized (GeneArt, Regensburg, Germany) and subcloned into a modified pFastBac vector, with the resulting construct encoding the GP67 secretion signal sequence at the N-terminus, and a His8 tag, followed by a TEV cleavage site at the C-terminus^32^. His8-tagged scFv16 was expressed and secreted by Sf9 insect cells, as previously reported^32^. Sf9 cells were collected by centrifugation at 5,000 ×g for 10 min, and the secreta-containing supernatant was combined with 5 mM CaCl_2_, 1 mM NiCl_2_, 20 mM HEPES, pH 8.0, and 150 mM NaCl. The supernatant was mixed with Ni Sepharose excel (Cytiva) and stirred for 1 h at 4°C. The bound resin was washed with buffer containing 20 mM HEPES, pH 8.0, 500 mM NaCl, and 20 mM imidazole, and further washed with 10 column volumes of buffer containing 20 mM HEPES, pH 8.0, 500 mM NaCl, and 20 mM imidazole. The protein was then eluted with 20 mM Tris, pH 8.0, 500 mM NaCl, and 400 mM imidazole, and the eluted fraction was concentrated and loaded onto a Superdex 200 10/300 Increase size-exclusion column, equilibrated in buffer containing 20 mM Tris (pH 8.0) and 150 mM NaCl. Peak fractions were pooled, concentrated to 5 mg ml^−1^ with a centrifugal filter device (10-kDa MW cut-off; MilliporeSigma, Burlington, MA, USA), and frozen in liquid nitrogen.

### Formation and purification of the GPR34-G_i_ complex

Purified GPR34-GFP was mixed with a 1.2 molar excess of G_i_ heterotrimer, scFv16, and TEV protease. After the addition of apyrase (to catalyse hydrolysis of unbound GDP) and agonist (final concentration, 10 μM), the coupling reaction was performed overnight at 4°C. To remove excess G protein, the complexing mixture was purified by M1 anti-Flag affinity chromatography. Bound complex was washed in buffer, containing 20 mM Tris HCl, pH 8.0, 150 mM NaCl, 0.01% GDN, 1 μM agonist, 10% glycerol, and 5 mM CaCl_2_. The complex was then eluted in 20 mM Tris-HCl, pH 8.0, 150 mM NaCl, 0.01% GDN, 10 μM agonist, 10% glycerol, 5 mM EDTA, and Flag peptide. The GPR34-G_i_-scFv16 complex was purified by size exclusion chromatography on a Superdex 200 10/300 column in 20 mM Tris-HCl, pH 8.0, 150 mM NaCl, 0.01% GDN, and 1 μM agonist, and peak fractions were concentrated to ~12 mg ml^−1^ for electron microscopy studies.

### Cryo-EM grid preparation and data collection

For cryo-EM grid preparation of GPR34-G_i_ complexes, 3 μl of protein at a concentration of approximately 10 mg ml^−1^ were loaded onto glow-discharged holey carbon grids (Quantifoil Au 300 mesh R1.2/1.3 or Quantifoil Cu/Rh 300 mesh R1.2/1.3), after which, these were plunge-frozen in liquid ethane, using a Vitrobot Mark IV (Thermo Fischer Scientific). Cryo-EM imaging was collected on a Titan Krios at 300 kV, using a Gatan K3 Summit detector. Images were obtained at a dose rate of about 8.0 e−/Å2 s−1, with a defocus ranging from −1.2 to −2.2 μm, using SerialEM software^46^. Total exposure time was 8 s, with 40 frames recorded per micrograph. A total of 2,358 and 2,674 videos were collected for S3E-LysoPS- and M1-bound GPR34-G_i_ complexes, respectively.

### Image processing

Single-particle analysis of GPR34-G_i_ complexes was performed with RELION-3.1^47,48^. Dose-fractionated image stacks were subjected to motion correction by MotionCorr2^49^, and contrast transfer function (CTF) parameters for micrographs were estimated by CTFFIND-4.0^50^.

For the S3E-LysoPS–GPR34-G_ii_ complex, a total of 2,012,061 particles were extracted,
 and the initial model was generated in RELION 3.1. Particles were subjected to several rounds of two-dimensional (2D) and three-dimensional (3D) classifications, resulting in the optimal classes of particles and yielding 258,700 particles. Particles were next subjected to 3D refinement, CTF refinement, and Bayesian polishing^51^. Following 3D refinement, particles were further classified into four classes, without alignment, using a mask covering the receptor. The 109,160 particles in the best class were subjected to 3D refinement and then further classified into three classes, without alignment, using a mask covering the extracellular half of the receptor. The 79,925 particles in the best class were subjected to 3D refinement, and postprocessing yielded a map having a nominal overall resolution of 3.4 Å, with the gold standard Fourier Shell Correlation (FSC=0.143) criteria^52^. The 3D model was further refined with a mask on the receptor, and as a result, the receptor has a higher nominal resolution of 3.6 Å. Local resolution was estimated by RELION 3.1. The processing strategy is described in Extended Data Figure 1.

For the M1-bound GPR34-G_i_ complex, a total of 1,823,992 particles were extracted, and the initial model was generated in RELION 3.1. Particles were subjected to several rounds of 2D and 3D classifications, resulting in the optimal classes of particles and yielding 485,175 particles. Particles were next subjected to 3D refinement, CTF refinement, and Bayesian polishing^51^.

Following 3D refinement, particles were further classified into four classes, without alignment, using a mask covering the extracellular half of the receptor. The 336,018 particles in the best class were subjected to 3D refinement and then further classified into four classes, without alignment, using a mask covering the complex. The 229,124 particles in the best class were subjected to 3D refinement, and postprocessing yielded a map having a nominal overall resolution of 3.3 Å, with the gold standard Fourier Shell Correlation (FSC=0.143) criteria. The 3D model was further refined with a mask on the receptor, and as a result, the receptor has a higher nominal resolution of 3.4 Å. Local resolution was estimated by RELION 3.1. The processing strategy is described in Extended Data Figure 1.

### Model building and refinement

The quality of the micelle-subtracted density map was sufficient to build a model manually in COOT^53,54^. Model building for the S3E-LysoPS bound GPR34-G_i_ complex was facilitated by the predicted GPR34 model in AlphaFold Protein Structure Database (https://alphafold.ebi.ac.uk/entry/Q9UPC5) and the cryo-EM structure of the μOR– G_i_ complex (PDB 6DDE). We manually modelled GPR34, the G_i_ heterotrimer, and scFv16 into the map by jiggle fit using COOT. The TM6 helix was manually fit into the density in COOT. We then manually readjusted the model into the density map using COOT and refined it using phenix.real_space_refine (v.1.19)^55,56^, with secondary-structure restraints imposed using phenix.secondary_structure_restraints. Finally, we refined the model using servalcat^57^. Model building for the M1-bound GPR34-G_i_ complex was initiated from the S3E-bound structure and followed the same procedure.

### Molecular dynamics (MD) simulations and docking simulation

The models of M1- and S3E-LysoPS-bound GPR34 were constructed based on the crystal structures and processed using Protein Preparation Wizard in Maestro2019-3 (Schrödinger, LLC, New York, NY, USA). The missing intracellular loop (ICL)2 of M1-bound GPR34 was modelled and refined using Prime. The N- and C-termini were omitted and capped with N-acetyl and N-methyl amide groups, respectively. Protonation states were optimized using PROPKA, and the whole structure of the ligand–receptor complex was minimized locally (force field, OPLS3e^58,59^) before the solvent model was added. Initial ligand–receptor complex models were embedded in a 1-palmitoyl-2-oleoyl-sn-glycero-3-phosphatidylcholine (POPC) membrane. The system was solvated with TIP3P water molecules and neutralized by adding 0.15 M NaCl. The prepared system contains ~40,000 atoms in total. The system was first subjected to preparation MD, followed by 1 μs of production MD simulation in the GPU Desmond suite (v.3.8.5.19)^60^. MD simulations were performed in the NPγT ensemble at 300 K using Langevin dynamics, and long-range electrostatic interactions were computed using the u-series algorithm^61^.

For *sn*-1 and *sn-*2 LysoPS-bound GPR34, the proper 3D conformation and ionization states of ligands (*sn*-1 and *sn-*2 LysoPS) generated using LigPrep, under the OPLS3e force field^59,58^, were used for ligand docking. The molecules were docked to the grid generated from the S3E-LysoPS-bound crystal structure, using the Glide SP mode, and strain correction was applied in the post-docking score. As a result, the docking pose with the best glide score was selected for each ligand and subjected to MD simulation, following the same protocol used for the M1- and S3E-LysoPS-bound structures, outlined above. The average structure of each complex was selected with the smallest root-mean-square deviation (RMSD) value from the average coordinates of ligand–protein complex atoms during 1-μs MD simulations.

### Supplementary information

#### Insight into ethoxy substitution

Intriguingly, although our structure–function relationship (SAR) results suggest enhanced activity by introducing an ethoxy group at the *sn*-1 position, the populated positions of the ethoxy substituent in M1 (alongside TM4) and S3E-LysoPS (pointing at TM5) are different (Fig, 5a,b, Extended Data Fig. 7a, and Extended Data Video 1). In both the cryo-EM structures and MD simulations, the ethoxy group does not make apparent interactions with receptor residues (Figs. 3e and 4b). Instead, this pinning ethoxy group tightly restricts and stabilizes the overall binding poses of the ligands (M1 and S3E-LysoPS), including both the hydrophilic moiety (root-mean-square Fluctuation [RMSF]_P_ <2 Å, RMSF_Ser_ <1 Å) and hydrophobic moiety (Extended Data Fig. 7b,c), by filling up the space in the vicinity of TMs 4–6 and extracellular loop (ECL)2.

#### Docking of *sn* −1 LysoPS derivative

In our previous studies of LysoPS derivatives, we could increase ligand activity of an *sn*-1-type derivative to a level comparable to that of *sn*-3 ligands by properly modifying the structure of the fatty acid moiety^16^ (Extended Data Fig. 7i). In this sense, the ligand’s hydrophobic tail can play a role similar to that of the ethoxy group in S3E-LysoPS, enabling the hydrophilic regions of *sn*-1-type ligands to adopt the proper U-shaped conformation required to interact with the binding site in a manner similar to *sn*-3 type derivatives (Extended Data Fig. 7i). This implies that structural change in the glycerol linker (*e.g*., S3E-LysoPS), as well as in the hydrophobic region, can induce the proper conformational changes in the phosphoserine part of the molecule, facilitating interaction with the binding pocket of GPR34, and thus activating GPR34 activity^62^.

#### Extended Data Video 1. 1-μs molecular dynamics (MD) simulation of GPR34 bound to the various agonists

The MD simulation movie was recorded every 600 ps. Hydrophilic network residues (*e.g*., K128^3.26^, F205^ECL2^, R286^6.55^, N309^7.35^, and E310^7.36^) are shown as sticks. The hydrophilic binding sites of GPR34 contain an assembly of charged amino acid residues that can efficiently interact with the charged phosphoserine. Therefore, hydrophilic interactions of the ligand in the binding pocket could be swapped. In fact, in the 1-μs MD simulation of *sn*-1 LysoPS, the *sn*-1-type hydrophilic interaction mode switches to the *sn*-3 type with a very low frequency. Conversely, the hydrophobic interactions are the same for *sn*-1, *sn*-2, and *sn*-3 LysoPS derivatives.

