## Supplementary file for "Structural basis for lysophosphatidylserine recognition by GPR34"

#### Extended Data Table 1. TGFα shedding assay.

Parameters obtained from TGFα shedding assay. The values of $pEC_{50}$, $EC_{50}$, and $E_{max}$, toward two agonists are shown.

| Compound | $pEC_{50}$ | $EC_{50}$ | $E_{max}$ | *n* = |
|---|---|---|---|---|
| *sn*-1 LysoPS (18:1) | 6.45 ± 0.25 | 350 nM | 7.5 ± 1.2% | 3 |
| *sn*-2 LysoPS (18:1) | 6.52 ± 0.09 | 300 nM | 13 ± 1.5% | 3 |
| S3E-LysoPS | 8.03 ± 0.02 | 9.4 nM | 11 ± 0.3% | 3 |
| M1 | 7.77 ± 0.01 | 17 nM | 14 ± 0.1% | 3 |

#### Extended Data Table 2. cAMP assay.

Parameters obtained from cAMP assay. The values of $pEC_{50}$, $EC_{50}$, and $E_{max}$, toward two agonists are shown.

| Compound | $pEC_{50}$ | $EC_{50}$ | $E_{max}$ | *n* = |
|---|---|---|---|---|
| *sn*-1 LysoPS (18:1) | < 5.5 | > 3 μM | NA | 3 |
| *sn*-2 LysoPS (18:1) | 6.85 ± 0.06 | 140 nM | 10 ± 1.3% | 3 |
| S3E-LysoPS | 8.07 ± 0.04 | 8.5 nM | 10 ± 0.7% | 3 |
| M1 | 7.80 ± 0.03 | 16 nM | 13 ± 0.4% | 3 |

#### Extended Data Table 3. Cryo-EM data collection, refinement, and validation statistics.

| | S3E-LysoPS-GPR34-$G_i$ | S3E-LysoPS-GPR34-$G_i$ (Receptor focused) | M1-GPR34-$G_i$ | M1-GPR34-$G_i$ (Receptor focused) |
|---|---|---|---|---|
| EMDB | EMD-34512 | EMD-34513 | EMD-34514 | EMD-34515 |
| PDB | 8H6W | 8H6X | 8H6Y | 8H6Z |
| **Data collection** | | | | |
| Microscope | Titan Krios (Thermo Fisher Scientific) | | | |
| Voltage (keV) | 300 | | | |
| Electron exposure (e–/Å$^2$) | 50 | | | |
| Detector | Gatan K3 summit camera (Gatan) | | | |
| Magnification | ×105,000 | | | |
| Defocus range (μm) | -0.8－-1.6 | | | |
| Pixel size (Å/pix) | 0.83 | | | |
| Number of movies | 2,358 | | 2,674 | |
| Symmetry | C1 | | | |
| Picked particles | 2,012,061 | | 1,823,992 | |
| Final particles | 79,925 | | 229,124 | |
| Map resolution (Å) | 3.4 | 3.6 | 3.3 | 3.4 |
| FSC threshold | 0.143 | | | |
| **Model refinement** | | | | |
| Atoms | 8,809 | 2,561 | 8,912 | 2,576 |
| R.m.s. deviations for ideal | | | | |
| Bond lengths (Å) | 0.013 | 0.012 | 0.013 | 0.0123 |
| Bond angles (°) | 1.357 | 1.31 | 1.449 | 1.3 |
| **Validation** | | | | |
| Clashscore | 10.19 | 10.53 | 9.1 | 12.57 |
| Rotamers (%) | 0 | 0 | 0.72 | 0 |
| Ramachandran plot | | | | |
| Favored (%) | 96.34 | 96.08 | 95.61 | 98.05 |
| Allowed (%) | 3.66 | 3.92 | 4.21 | 1.95 |
| Outlier (%) | 0 | 0 | 0.18 | 0 |

### Extended Data Table 4. Mutagenesis analysis of GPR34.

Parameters obtained from the TGFα shedding assay. The values of $pEC_{50}$, $EC_{50}$, $E_{max}$, and $log_{10}$ relative intrinsic activity (RAi) toward two agonists are shown with the $log_{10}$ mean fluorescence intensity (MFI), indicating the cell surface expression level of each receptor.

| Construct | | Ligand response | | | | | | | | | | Surface expression | |
|---|---|---|---|---|---|---|---|---|---|---|---|---|---|
| | | S3E-LysoPS | | | | | M1 | | | | | | |
| Mutation | BW position | $pEC_{50}$ | $EC_{50}$ | $E_{max}$ | LogRAi | *n* = | $pEC_{50}$ | $EC_{50}$ | $E_{max}$ | LogRAi | *n* = | LogMFI | *n* = |
| WT | | 8.02 ± 0.04 | 9.6 nM | 10 ± 1% | 0 | 8 | 7.86 ± 0.01 | 14 nM | 13 ± 1.2% | 0 | 8 | 2.69 ± 0.05 | 10 |
| R110A | 2.6 | 7.88 ± 0.12 | 13 nM | 7.7 ± 1.7% | -0.25 ± 0.11 | 5 | 7.68 ± 0.13 | 21 nM | 7 ± 1.5% | -0.49 ± 0.13 | 5 | 2.49 ± 0.03 | 5 |
| K128A | 3.26 | 8.00 ± 0.09 | 10 nM | 7.4 ± 1.8% | -0.16 ± 0.1 | 5 | 6.35 ± 0.19 | 450 nM | 13 ± 2% | -1.54 ± 0.15 | 5 | 2.59 ± 0.06 | 5 |
| Y135A | 3.33 | < 5.5 | > 3 μM | < 2% | < -3 | 5 | < 5.5 | > 3 μM | < 2% | < -3 | 5 | 2.61 ± 0.07 | 5 |
| Y135F | 3.33 | 7.74 ± 0.04 | 18 nM | 8.5 ± 1.9% | -0.36 ± 0.06 | 5 | 7.36 ± 0.05 | 43 nM | 7.3 ± 1.6% | -0.79 ± 0.05 | 5 | 2.55 ± 0.05 | 5 |
| M136A | 3.34 | 8.08 ± 0.08 | 8.2 nM | 6.1 ± 1.2% | -0.15 ± 0.08 | 5 | 7.71 ± 0.03 | 20 nM | 10 ± 1.5% | -0.27 ± 0.03 | 5 | 2.61 ± 0.06 | 5 |
| M189A | 4.6 | 7.65 ± 0.08 | 22 nM | 9.1 ± 1.5% | -0.40 ± 0.08 | 5 | 8.23 ± 0.04 | 5.9 nM | 14 ± 1.7% | 0.40 ± 0.05 | 5 | 2.63 ± 0.07 | 5 |
| F205A | ECL2 | < 5.5 | > 3 μM | < 2% | < -3 | 5 | < 5.5 | > 3 μM | < 2% | < -3 | 5 | 2.47 ± 0.06 | 5 |
| H206A | ECL2 | 7.47 ± 0.12 | 34 nM | 17 ± 3.5% | -0.34 ± 0.13 | 5 | 6.83 ± 0.09 | 150 nM | 25 ± 3.8% | -0.78 ± 0.08 | 5 | 2.60 ± 0.07 | 5 |
| Y207A | ECL2 | 6.07 ± 0.04 | 850 nM | 15 ± 3.3% | -1.77 ± 0.04 | 5 | 6.21 ± 0.03 | 610 nM | 14 ± 3% | -1.65 ± 0.05 | 5 | 2.66 ± 0.09 | 5 |
| R208A | ECL2 | 6.68 ± 0.26 | 210 nM | 12 ± 3.2% | -1.29 ± 0.21 | 5 | 6.71 ± 0.10 | 190 nM | 14 ± 2.2% | -1.15 ± 0.08 | 5 | 2.57 ± 0.05 | 5 |
| K210A | ECL2 | 8.30 ± 0.11 | 5 nM | 15 ± 2% | 0.45 ± 0.11 | 5 | 7.29 ± 0.06 | 51 nM | 18 ± 3.3% | -0.45 ± 0.07 | 5 | 2.64 ± 0.04 | 5 |
| F219A | 5.39 | 7.52 ± 0.08 | 30 nM | 16 ± 2.6% | -0.28 ± 0.07 | 5 | 8.36 ± 0.04 | 4.3 nM | 22 ± 2.8% | 0.71 ± 0.05 | 5 | 2.69 ± 0.07 | 5 |
| N220A | 5.4 | 7.71 ± 0.10 | 19 nM | 14 ± 2.1% | -0.15 ± 0.09 | 5 | 6.87 ± 0.09 | 140 nM | 18 ± 3.9% | -0.89 ± 0.08 | 5 | 2.55 ± 0.04 | 5 |
| L223A | 5.43 | 7.41 ± 0.27 | 39 nM | 14 ± 2.7% | -0.47 ± 0.23 | 5 | 6.84 ± 0.06 | 140 nM | 16 ± 2.8% | -0.96 ± 0.07 | 5 | 2.52 ± 0.05 | 5 |
| R286A | 6.55 | < 5.5 | > 3 μM | < 2% | < -3 | 5 | < 5.5 | > 3 μM | < 2% | < -3 | 5 | 2.55 ± 0.04 | 5 |
| Y289A | 6.58 | < 5.5 | > 3 μM | < 2% | < -3 | 5 | < 5.5 | > 3 μM | < 2% | < -3 | 5 | 2.50 ± 0.03 | 5 |
| N309A | 7.35 | 6.87 ± 0.09 | 130 nM | 12 ± 2.9% | -1.09 ± 0.09 | 5 | 7.06 ± 0.05 | 87 nM | 11 ± 2.4% | -0.91 ± 0.07 | 5 | 2.48 ± 0.04 | 5 |
| E310A | 7.36 | 6.50 ± 0.11 | 320 nM | 9.1 ± 2.6% | -1.58 ± 0.10 | 5 | 6.70 ± 0.07 | 200 nM | 7.6 ± 1.8% | -1.44 ± 0.10 | 5 | 2.45 ± 0.04 | 5 |

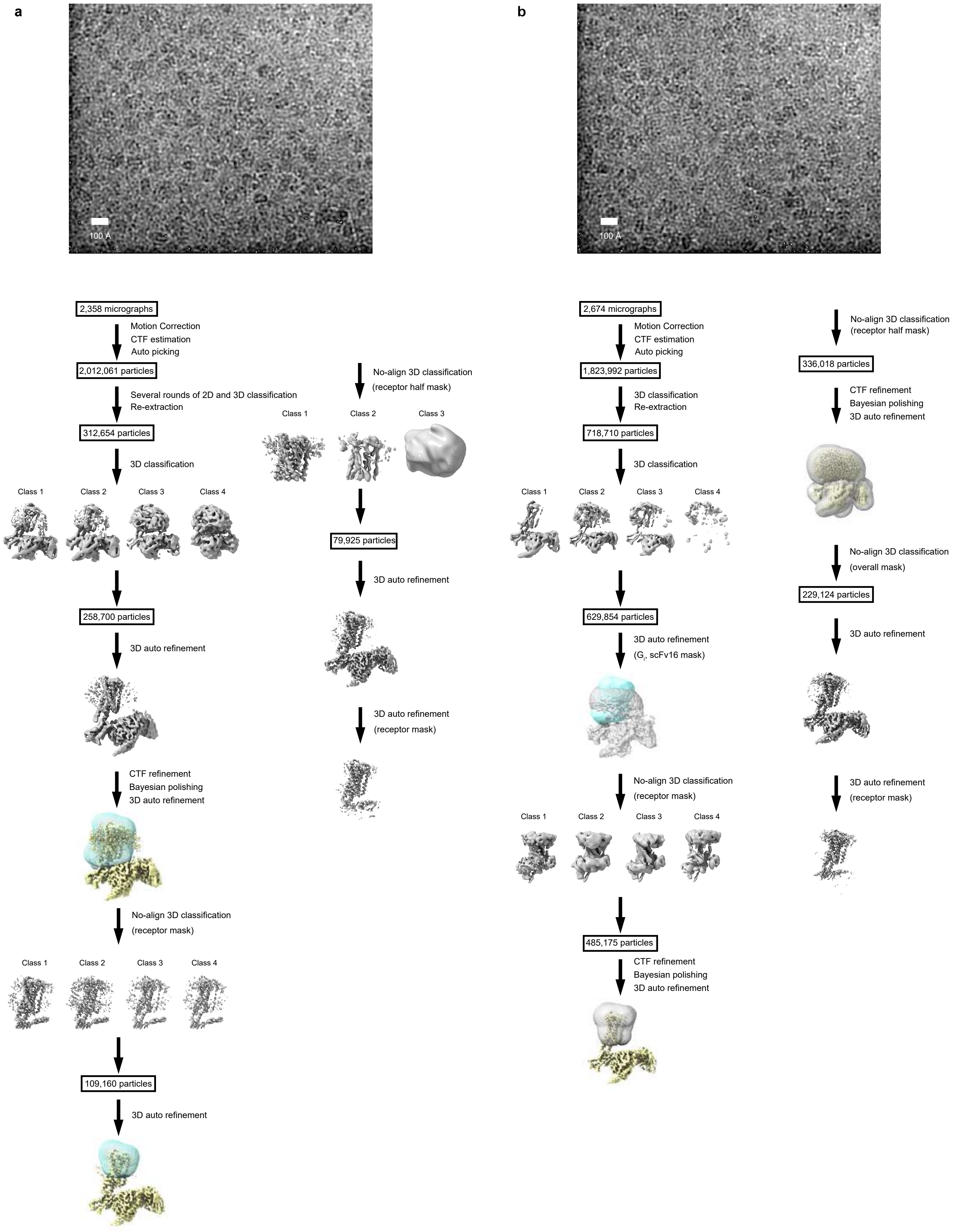


#### Extended Data Fig. 1. Cryo-EM workflows of the GPR34-$G_i$ complexes.

Flow chart of the cryo-EM data processing for the GPR34-$G_i$ complexes bound to S3E-LysoPS and M1, including particle projection selection, classification, and 3D density map reconstruction. Details are provided in the Methods section.

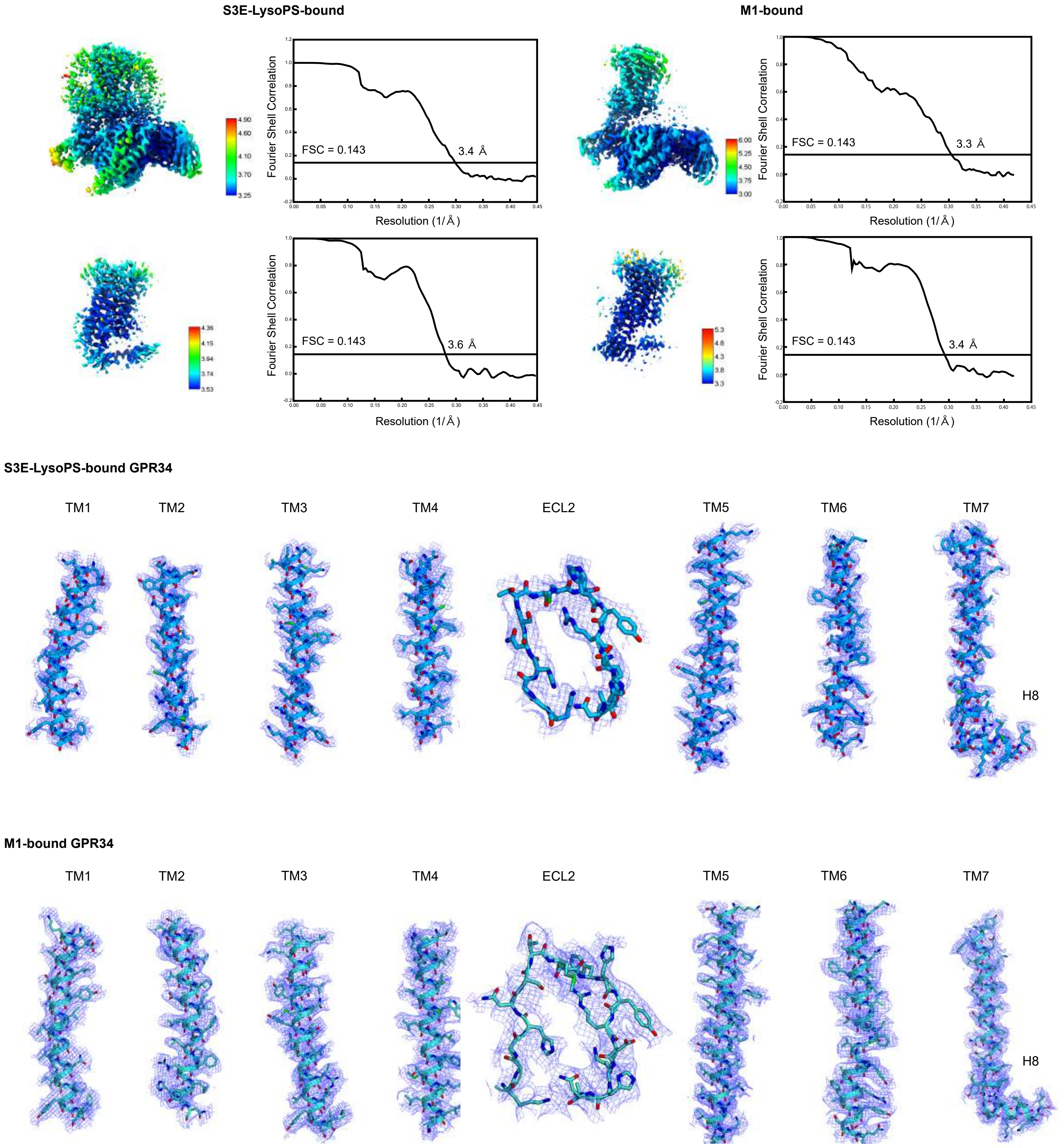


### Extended Data Fig. 2. Maps and model quality.

Local resolution maps, FSC curves, and cryo-EM density maps of the GPR34-$G_i$ complexes bound to S3E-LysoPS and M1.

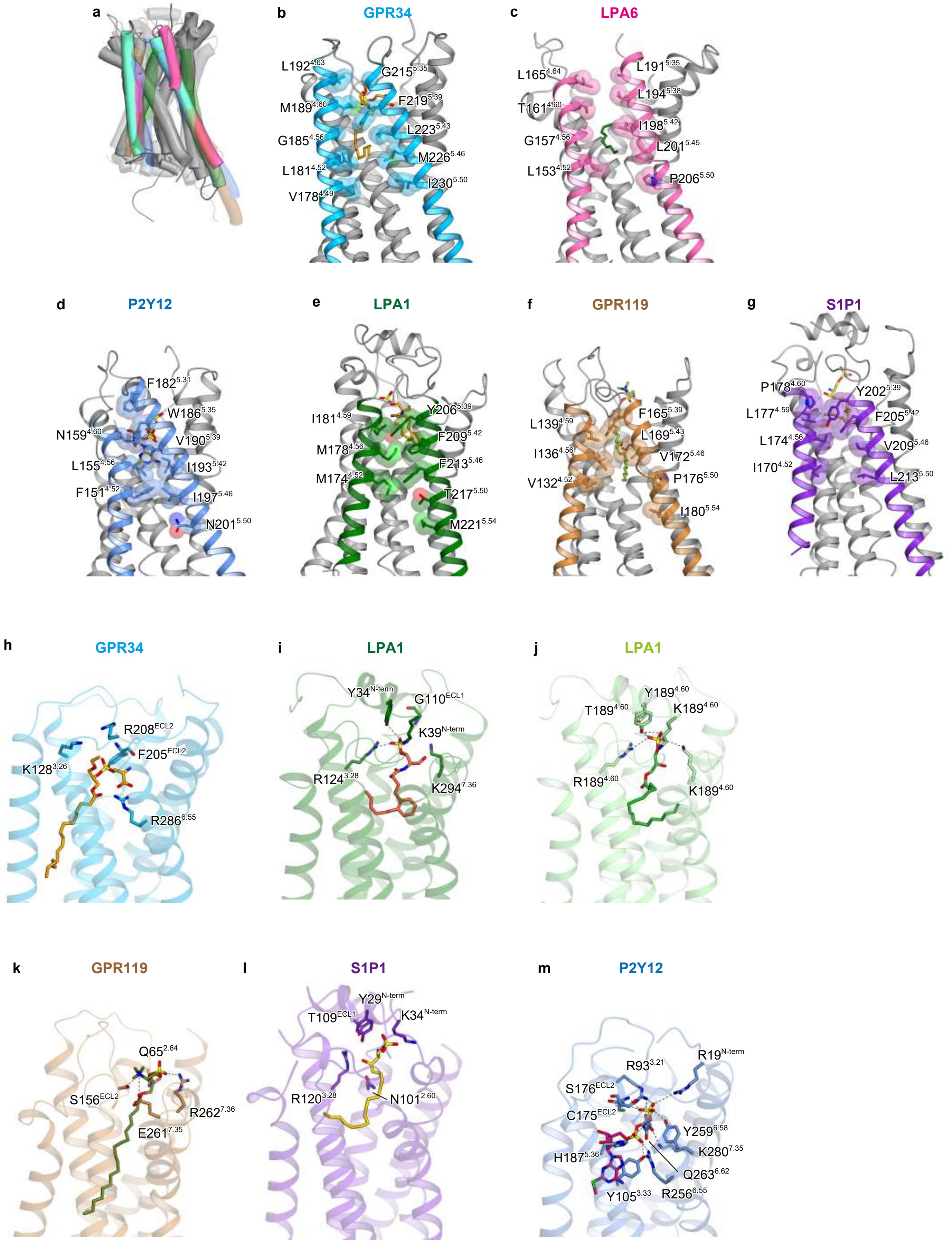


#### Extended Data Fig. 3. Structural comparison of GPR34 with other GPCRs.

**(a)** Superimposition of the structures of GPR34, $LPA_6$, $LPA_1$, GPR119, and $S1P_1$.

**(b-g)** Interactions between the extracellular halves of TM4 and TM5 in respective receptors.

**(h-m)** Comparison of the agonist binding modes in respective receptors.

a

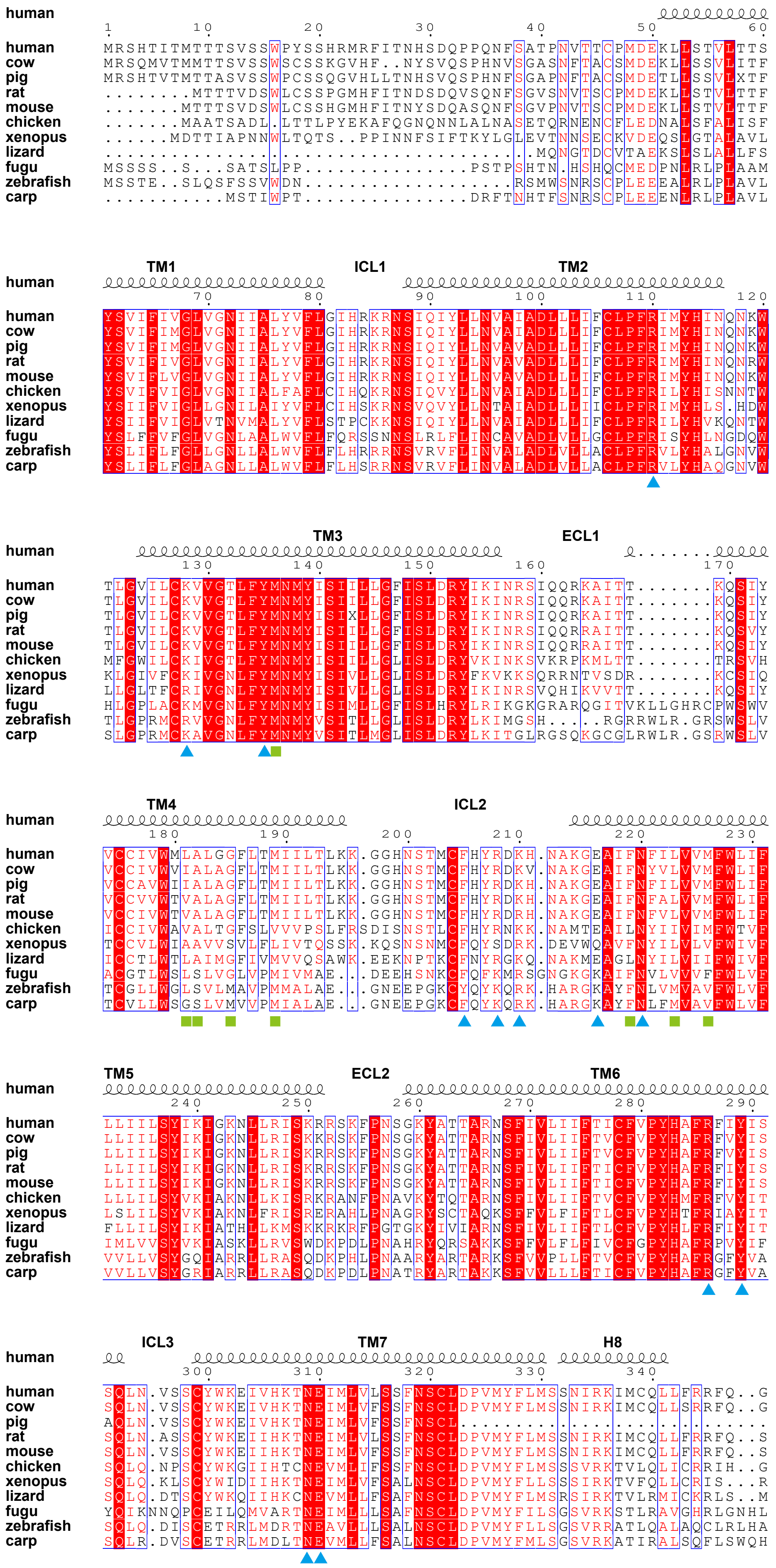

human
1 10 20 30 40 50 60
human MRSHTITMTTTSVSSWPYSSHRMRFITNHSDQPPQNFSATPNVTTCPMDEKLLSTVLTTS
cow MRSQMVTMMTTSVSSWSCSSKGVHF..NYSVQSPHNVSGASNFTACSMDEKLLSSVLITF
pig MRSHTVTMTTASVSSWPCSSQGVHLLTNHSVQSPHNFSGAPNFTACSMDETLLSSVLXTF
rat ........MTTTVDSWLCSSPGMHFITNDSDQVSQNFSGVSNVTSCPMDEKLLSTVLTTF
mouse .......MTTTSVDSWLCSSHGMHFITNYSDQASQNFSGVPNVTSCPMDEKLLSTVLTTF
chicken .......MAATSADL.LTTLPYEKAFQGNQNNLALNASETQRNENCFLEDNALSFALISF
xenopus ......MDTTIAPNNWLTQTS..PPINNFSIFTKYLGLEVTNNSECKVDEQSLGTALAVL
lizard .......................................MQNGTDCVTAEKSLSLALLFS
fugu MSSSS..S...SATSLPP...............PSTPSHTN.HSHQCMEDPNLRLPLAAM
zebrafish MSSTE..SLQSFSSVWDN...................RSMWSNRSCPLEEEALRLPLAVL
carp ...........MSTIWPT...............DRFTNHTFSNRSCPLEEENLRLPLAVL
TM1 ICL1 TM2
human
70 80 90 100 110 120
human YSVIFIVGLVGNIIALYVFLGIHRKRNSIQIYLLNVAIADLLLIFCLPFRIMYHINQNKW
cow YSVIFIMGLVGNIIALYVFLGIHRKRNSIQIYLLNVAIADLLLIFCLPFRIMYHINQNKW
pig YSVIFIMGLVGNIIALYVFLGIHRKRNSIQIYLLNVAVADLLLIFCLPFRIMYHINQNKW
rat YSVIFIVGLVGNIIALYVFLGIHRKRNSIQIYLLNVAVADLLLIFCLPFRIMYHINQNRW
mouse YSVIFLVGLVGNIIALYVFLGIHRKRNSIQIYLLNVAVADLLLIFCLPFRIMYHINQNKW
chicken YSVIFVIGLVGNIIALFAFLCIHQKRNSIQVYLLNVAIADLLLIFCLPFRILYHISNNTW
xenopus YSIIFVIGLLGNILAIYVFLCIHSKRNSVQVYLLNTAIADLLLIICLPFRIMYHLS.HDW
lizard YSIIFVIGLVTNVMALYVFLSTPCKKNSIQIYLLNVAIADLLLIFCLPFRILYHVKQNTW
fugu YSLFFVFGLVGNLAALWVFLFQRSSNNSLRLFLINCAVADLVLLGCLPFRISYHLNGDQW
zebrafish YSLIFLFGLLGNLLALWVFLFLHRRRNSVRVFLINVALADLVLLACLPFRVLYHALGNVW
carp YSLIFLFGLAGNLLALWVFLFLHSRRNSVRVFLINVALADLVLLACLPFRVLYHAQGNVW
TM3 ECL1
human
130 140 150 160 170
human TLGVILCKVVGTLFYMNMYISIILLGFISLDRYIKINRSIQQRKAITT.......KQSIY
cow TLGVILCKVVGTLFYMNMYISIILLGFISLDRYIKINRSIQQRKAITT.......KQSIY
pig TLGVILCKVVGTLFYMNMYISIXLLGFISLDRYIKINRSIQQRKAITT.......KQSIY
rat TLGVILCKVVGTLFYMNMYISIILLGFISLDRYIKINRSIQQRRAITT.......KQSVY
mouse TLGVILCKVVGTLFYMNMYISIILLGFISLDRYIKINRSIQQRRAITT.......KQSIY
chicken MFGWILCKIVGTLFYMNMYISIILLGLISLDRYVKINKSVKRPKMLTT.......TRSVH
xenopus KLGIVFCKIVGNLFYMNMYISIVLLGLISLDRYFKVKKSQRRNTVSDR.......KCSIQ
lizard LLGLTFCRIVGNLFYMNMYISIVLLGLISLDRYIKINRSVQHIKVVTT.......KQSIY
fugu HLGPLACKMVGNLFYMNMYISIMLLGFISLHRYLRIKGKGRARQGITVKLLGHRCPWSWV
zebrafish TLGPRMCRVVGNLFYMNMYVSITLLGLISLDRYLKIMGSH.....RGRRWLR.GRSWSLV
carp SLGPRMCKAVGNLFYMNMYVSITLMGLISLDRYLKITGLRGSQKGCGLRWLR.GSRWSLV
TM4 ICL2
human
180 190 200 210 220 230
human VCCIVWMLALGGFLTMIILTLKK.GGHNSTMCFHYRDKH.NAKGEAIFNFILVVMFWLIF
cow VCCIVWVIALAGFLTMIILTLKK.GGHNSTMCFHYRDKV.NAKGEAIFNYVLVVMFWLIF
pig VCCAVWIIALAGFLTMIILTLKK.GGHNSTMCFHYRDKH.NAKGEAIFNFILVVMFWLIF
rat VCCVVWTVALAGFLTMIILTLKK.GGHNSTMCFHYRDKH.NAKGEAIFNFALVVMFWLIF
mouse VCCIVWTVALAGFLTMIILTLKK.GGHNSTMCFHYRDRH.NAKGEAIFNFVLVVMFWLIF
chicken ICCIVWAVALTGFSLVVVPSLFRSDISNSTLCFHYRNKK.NAMTEAILNYIIVIMFWTVF
xenopus TCCVLWIAAVVSVLFLIVTQSSK.KQSNSNMCFQYSDRK.DEVWQAVFNYILVLVFWIVF
lizard ICCTLWTLAIMGFIVMVVQSAWK.EEKNPTKCFNYRGKQ.NAKMEAGLNYILVIIFWIVF
fugu ACGTLWSLSLVGLVPMIVMAE...DEEHSNKCFQFKMRSGNGKGKAIFNVLVVVFFWLVF
zebrafish TCGLLWGLSVLMAVPMMALAE...GNEEPGKCYQYKQRK.HARGKAYFNLVMVAVFWLVF
carp TCVLLWSGSLVMVVPMIALAE...GNEEPGKCFQYKQRK.HARGKAYFNLFMVAVFWLVF
TM5 ECL2 TM6
human
240 250 260 270 280 290
human LLIILSYIKIGKNLLRISKRRSKFPNSGKYATTARNSFIVLIIFTICFVPYHAFRFIYIS
cow LLIILSYIKIGKNLLRISKKRSKFPNSGKYATTARNSFIVLIIFTVCFVPYHAFRFVYIS
pig LLIILSYIKIGKNLLRISKRRSKFPNSGKYATTARNSFIVLIIFTVCFVPYHAFRFVYIS
rat LLIILSYIKIGKNLLRISKRRSKFPNSGKYATTARNSFIVLIIFTICFVPYHAFRFIYIS
mouse LLIILSYIKIGKNLLRISKRRSKFPNSGKYATTARNSFIVLIIFTICFVPYHAFRFIYIS
chicken LLLILSYVKIAKNLLKISRKRANFPNAVKYTQTARNSFIVLIIFTVCFVPYHMFRFVYIT
xenopus LSLILSYVKIAKNLFRISRERAHLPNAGRYSCTAQKSFFVLFIFTLCFVPYHTFRIAYIT
lizard FLLILSYIKIATHLLKMSKKRKRFPGTGKYIVIARNSFIVLIIFTLCFVPYHLFRFIYIT
fugu IMLVVSYVKIASKLLRVSWDKPDLPNAHRYQRSAKKSFFVLFLFIVCFGPYHAFRPVYIF
zebrafish VVLLVSYGQIARRLLRASQDKPHLPNAARYARTARKSFVVPLLFTVCFVPYHAFRGFYVA
carp VVLLVSYGRIARRLLRASQDKPDLPNATRYARTAKKSFVVLLLFTICFVPYHAFRGFYVA
ICL3 TM7 H8
human
300 310 320 330 340
human SQLN.VSSCYWKEIVHKTNEIMLVLSSFNSCLDPVMYFLMSSNIRKIMCQLLFRRFQ..G
cow SQLN.VSSCYWKEIVHKTNEIMLVFSSFNSCLDPVMYFLMSSNIRKIMCQLLSRRFQ..G
pig AQLN.VSSCYWKEIVHKTNEIMLVFSSFNSCL............................
rat SQLN.ASSCYWKEIIHKTNEIMLVLSSFNSCLDPVMYFLMSSNIRKIMCQLLFRRFQ..S
mouse SQLN.VSSCYWKEIIHKTNEIMLVFSSFNSCLDPVMYFLMSSNIRKIMCQLLFRRFQ..S
chicken SQLQ.NPSCYWKGIIHTCNEVMLIFSSFNSCLDPVMYFLMSSSVRKTVLQLICRRIH..G
xenopus SQLQ.KLSCYWIDIIHKTNEIMLVFSALNSCLDPVMYFLLSSSIRKTVFQLLCRIS...R
lizard SQLQ.DTSCYWKQIIHKCNEVMLLFSAFNSCLDPVMYFLMSRSIRKTVLRMICKRLS..M
fugu YQIKNNQPCEILQMVARTNEIMLLLSAFNSCLDPVMYFILSGSVRKSTLRAVGHRLGNHL
zebrafish SQLQ.DISCETRRLMDRTNEAVLLLSALNSCLDPVMYFLLSGSVRRATLQALAQCLRLHA
carp SQLR.DVSCETRRLMDLTNEVMLLFSALNSCLDPVMYFMLSGSVRKATIRALSQFLSWQH

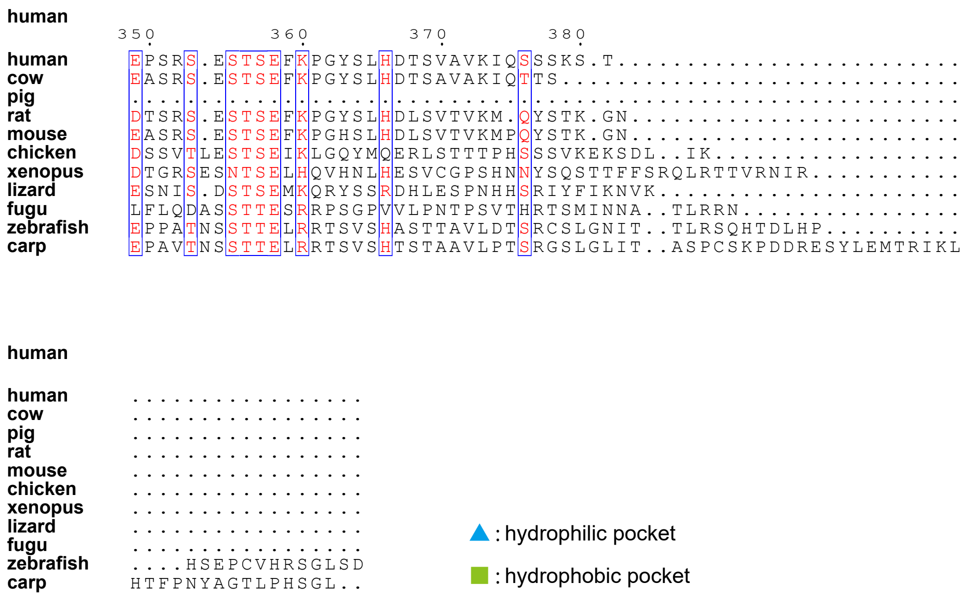


**b**

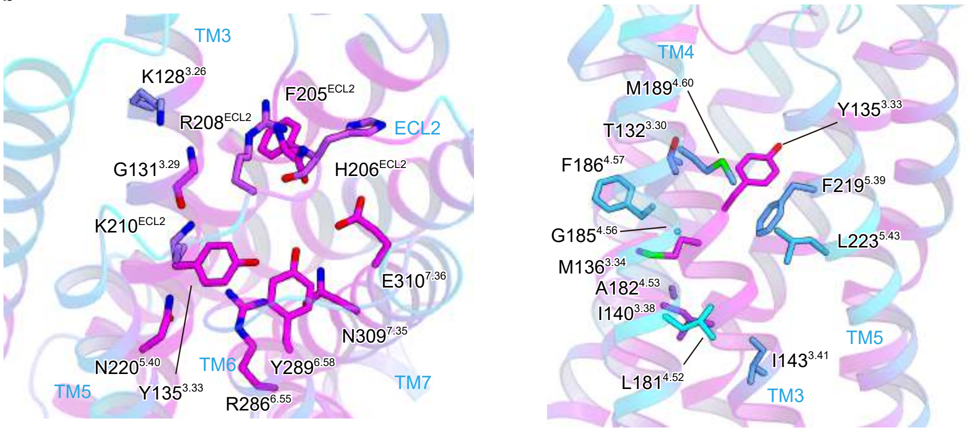


#### Extended Data Fig. 4. Alignment of GPR34 across vertebrates.

**(a)** Alignment of GPR34 across vertebrates. Amino acid sequence alignment of GPR34 from human (Uniprot ID: P0DMS8), cow (Uniprot ID: Q0VC81), pig (Uniprot ID: F1SBN8), rat (Uniprot ID: P28647), mouse (Uniprot ID: Q3U4C5), chicken (Uniprot ID: R4GIJ8), Xenopus (Uniprot ID: ENSXETT00000046736.1), lizard (Uniprot ID: G1KF19), and fugu (Uniprot ID: H2VBI6). Secondary structure elements for α-helices are indicated by cylinders. Conservation of the residues is indicated as follows: red panels for completely conserved; red letters for partially conserved; and black letters for not conserved.

**(b)** Conservation of the residues of GPR34. The sequence conservation among the vertebrate homologues of GPR34 was calculated using the ConSurf server (http://consurf.tau.ac.il) and is colored from cyan (low) to maroon (high).

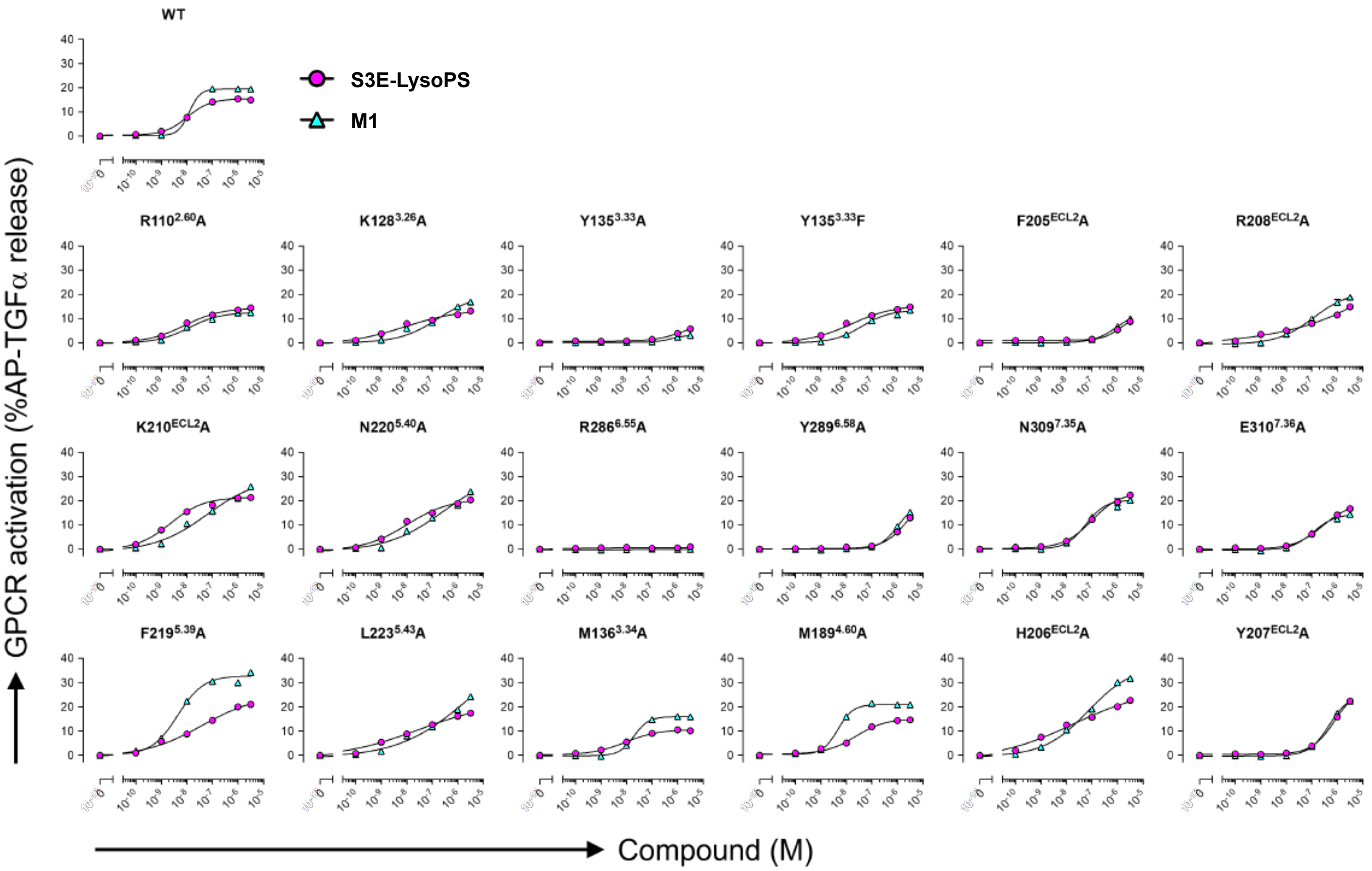


#### Extended Data Fig. 5. Mutational analyses using the TGFα shedding assay.

The responses of wild-type GPR34 and 18 mutants of GPR34 toward S3E-LysoPS and M1 are shown. The TGFα shedding assay was performed using HEK293 cells transfected with AP-TGFα, $G\alpha_{q/i1}$, and each GPR34-expressing vector. For negative control cells, empty plasmid was transfected instead of a receptor-expressing plasmid. In each panel, receptor-specific AP-TGFα release levels (differences in AP-TGFα release between receptor-expressing cells and negative control cells) are shown. Symbols and error bar mean average and SEM, respectively, from 5-8 independent experiments. See Table S4 for parameters obtained from the concentration-responses curves.

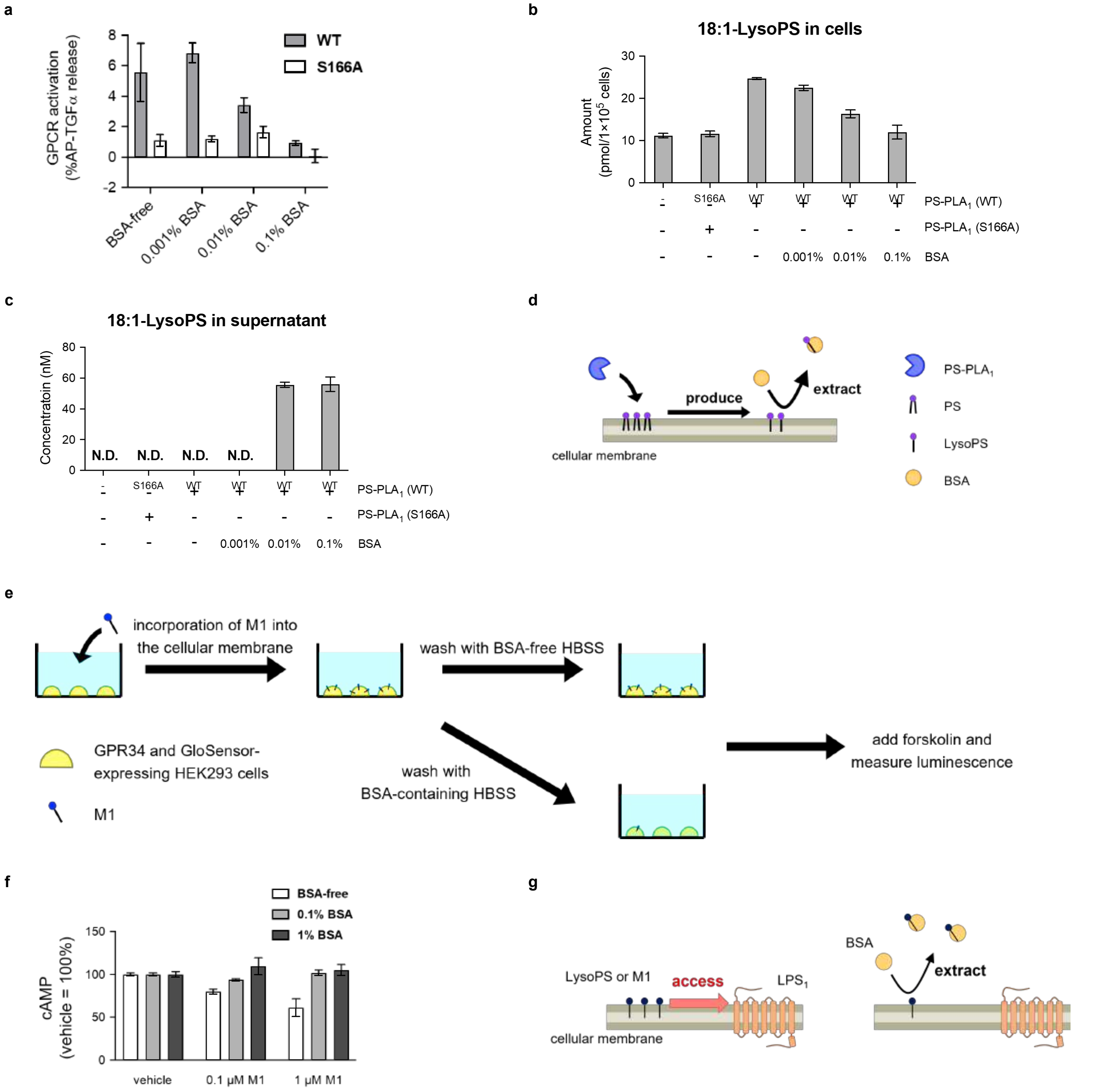


**Extended Data Fig. 6. LysoPS and M1 access GPR34 from its lateral side.**

**(a)** GPR34 activation induced by recombinant PS-PLA$_1$ and the effect of LysoPS extraction from the cellular membrane on it. HEK293A cells expressing AP-TGFα, Gα$_{q/i1}$, and GPR34 were stimulated by recombinant PS-PLA$_1$ with or without BSA. For the negative control, HEK293 cells transfected with empty plasmid instead of GPR34-encoding plasmid were used. After 1 hour incubation, the amount of AP-TGFα in the supernatant was determined. The receptor-specific AP-TGFα release (differences in AP-TGFα release between GPR34-expressing cells and negative control cells) are shown. S166A is a mutant PS-PLA$_1$, which do not have the catalytic activity. **(b and c)** The amount of 18:1-LysoPS in the cells treated with recombinant PS-PLA$_1$ and the concentration of 18:1-LysoPS in the supernatant. HEK293A cells were treated with recombinant PS-PLA$_1$ for 1 hour. Then, LysoPS in the cells and the supernatant was extracted into MeOH, and the amount was evaluated by LC-MS/MS. **(d)** The illustration of GPR34 activation induced by PS-PLA$_1$. PS-PLA$_1$ hydrolyzes phosphatidylserine in the cellular membrane to produce LysoPS. The LysoPS laterally spreads in the cellular membrane and accesses GPR34. **(e and f)** The effect of M1 incorporation into and extraction from the cellular membrane on GPR34 activation are shown. The schematic illustration of experimental procedure is shown (e). HEK293 cells expressing GPR34 and GloSensor-22F were incubated with M1 and washed by BSA-free or BSA-containing HBSS. After that, forskolin was added and the luminescence was measured. The cAMP levels in the HEK293 cells-expressing are shown (f). **(g)** The schematic illustration of M1 accessing GPR34. GPR34 can be activated by M1, which is in the cellular membrane. This is indicated by the data (e), which showed GPR34 was not activated when the cells were washed by BSA-containing HBSS and M1 was extracted from the cellular membrane.

a

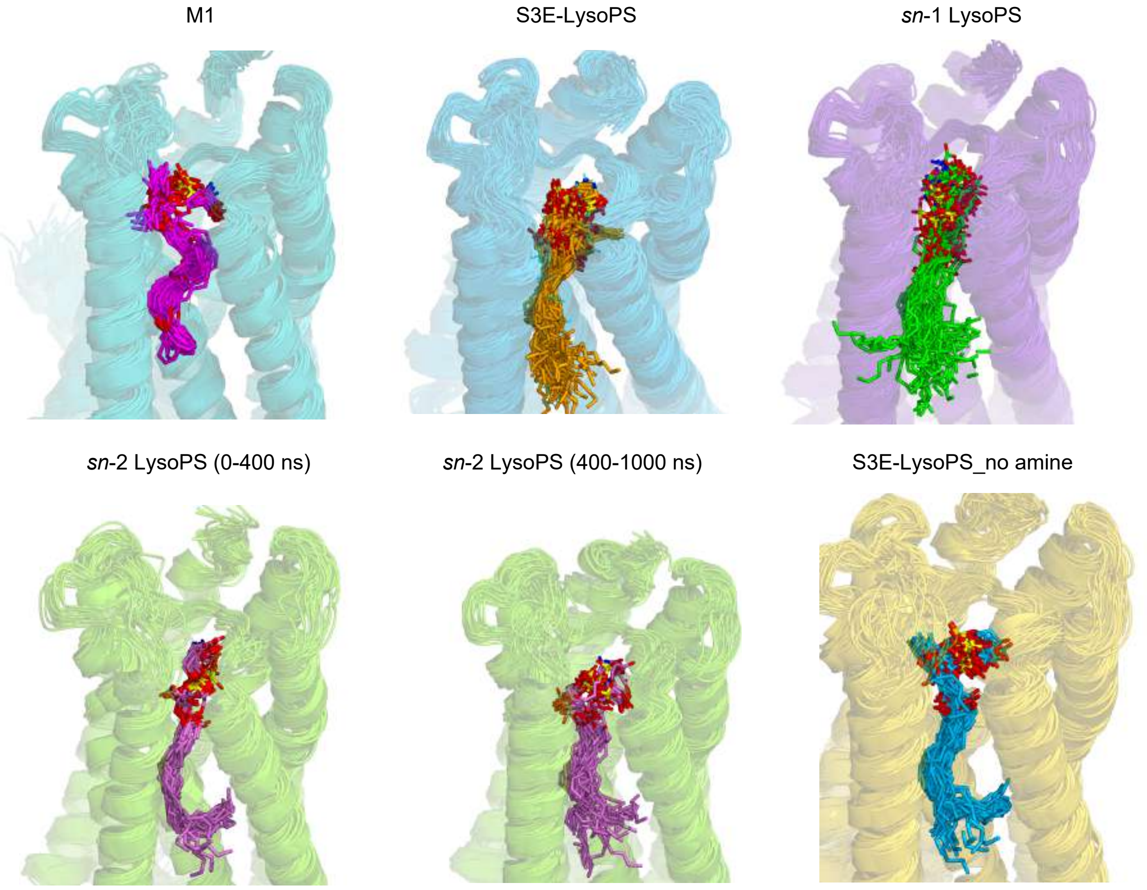
M1
S3E-LysoPS
*sn*-1 LysoPS
*sn*-2 LysoPS (0-400 ns)
*sn*-2 LysoPS (400-1000 ns)
S3E-LysoPS_no amine

b

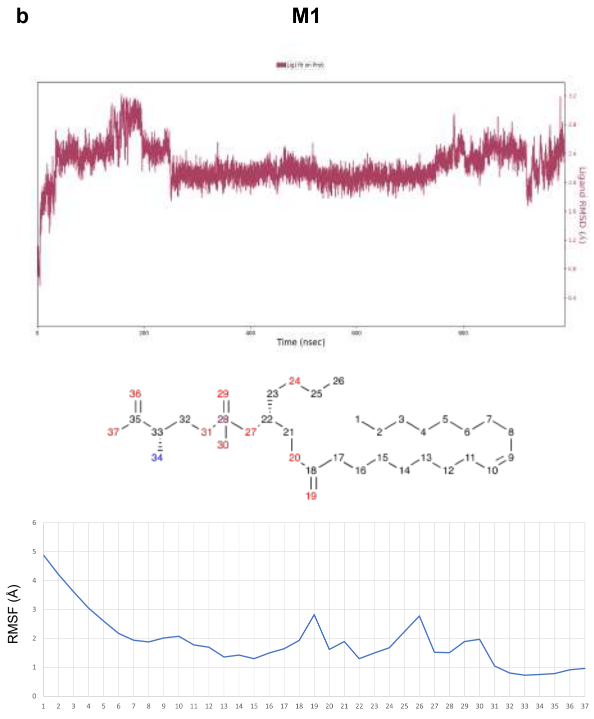
M1
Time (nsec)
Ligand RMSD (Å)
RMSF (A)

c

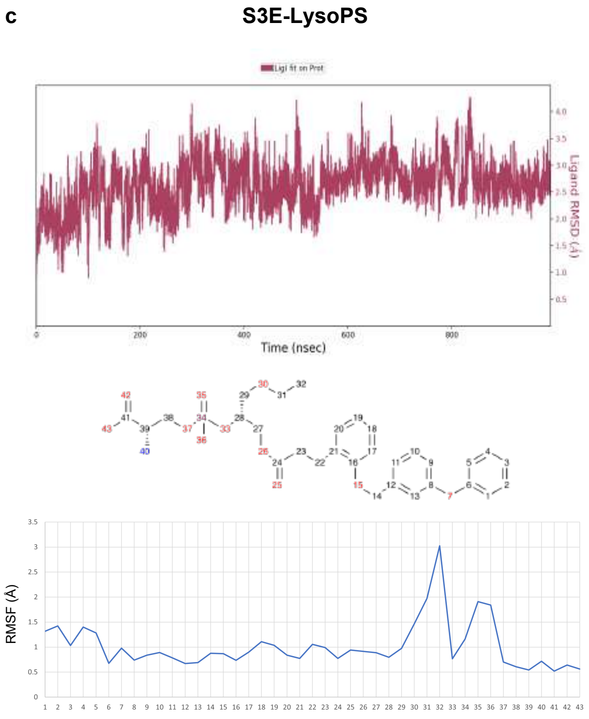
S3E-LysoPS
Lig1 fit on Prot
Time (nsec)
Ligand RMSD (Å)
RMSF (A)

d

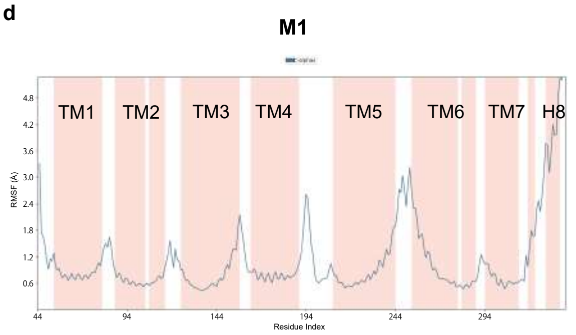
M1
TM1
TM2
TM3
TM4
TM5
TM6
TM7
H8
RMSF (A)
Residue Index

e

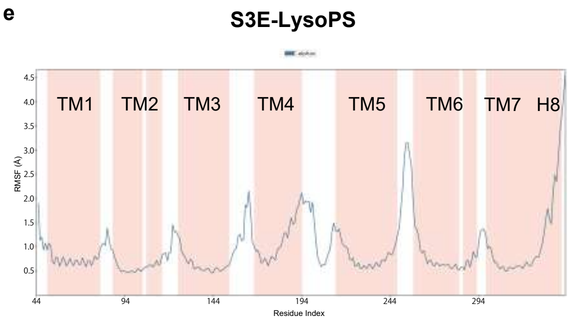
S3E-LysoPS
TM1
TM2
TM3
TM4
TM5
TM6
TM7
H8
RMSF (A)
Residue Index

**f** M1

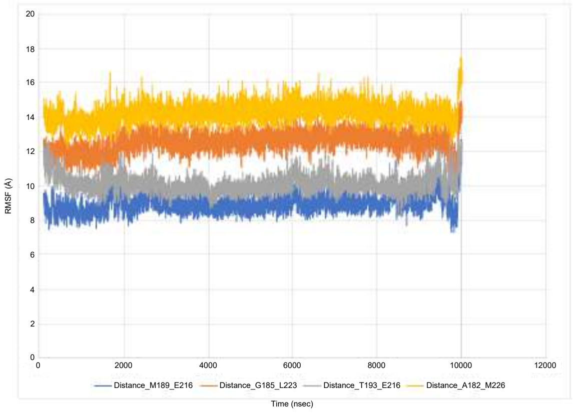


**g** S3E-LysoPS

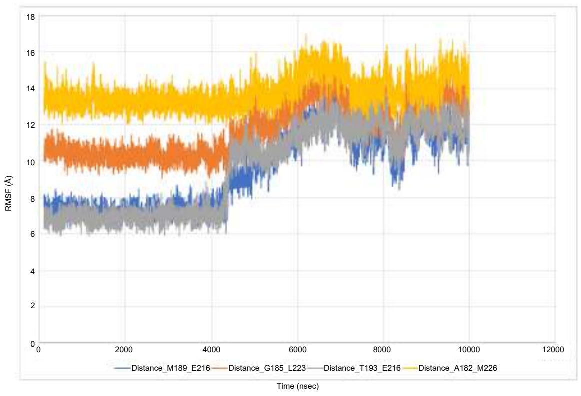


**h**

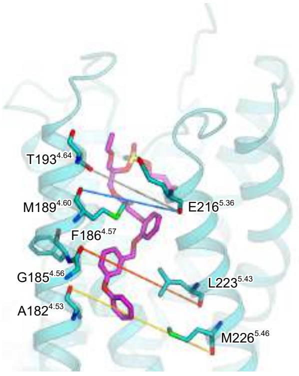


**i** *sn*-1 derivative
(deoxysn1_C3_Ph_o_OBn_m_OPh)

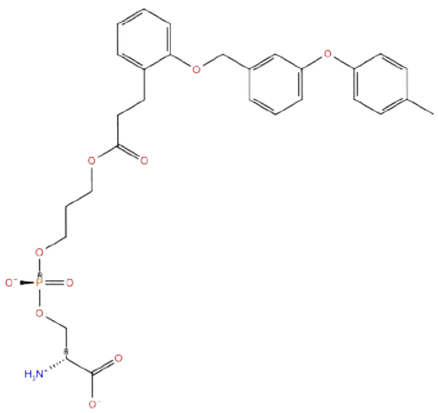

**j** M1-GPR34 (MD) *sn*-1 derivative-GPR34 (docking)

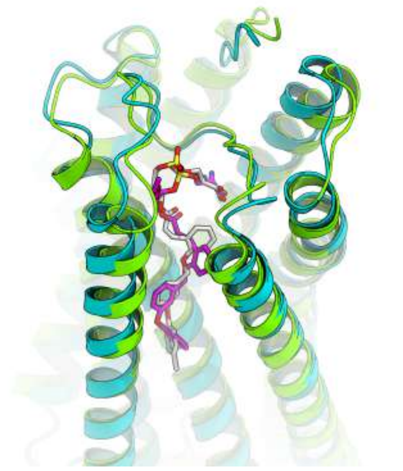

#### Extended Data Fig. 7. Molecular dynamics simulations.

**(a)** Ensembles from each trajectory. The complex structures were extracted every 20 ns and superimposed on the first structure. **(b and c)** Ligand RMSD (top panel) and RMSF values (bottom panel) during the 1 μs MD simulation. Atom index is illustrated on the chemical structure. **(d and e)** Protein RMSF values during the 1 μs MD simulation. **(f-h)** Distances between the residues in TM4 and TM5 during the MD simulations. The distances of M189$^{4.60}$-E216$^{5.36}$ G185$^{4.56}$-L223$^{5.43}$ T193$^{4.64}$-L223$^{5.43}$ A182$^{4.53}$-M226$^{5.46}$ were measured, and their positions in the average structure of the M1-bound receptor are shown in (H). **(i)** Chemical structure of the sn-1 derivative (deoxysn1_C3_Ph_o_OBn_m_OPh) mentioned in the paper. Its EC50 value for the human GPR34 was reportedly 27 nM in the TGFα shedding assay(*17*), and comparable to that of S3E-LysoPS. **(j)** Superimposition of the *sn*-1 derivative-docked structure and the average structure of the M1-bound receptor during the simulation.

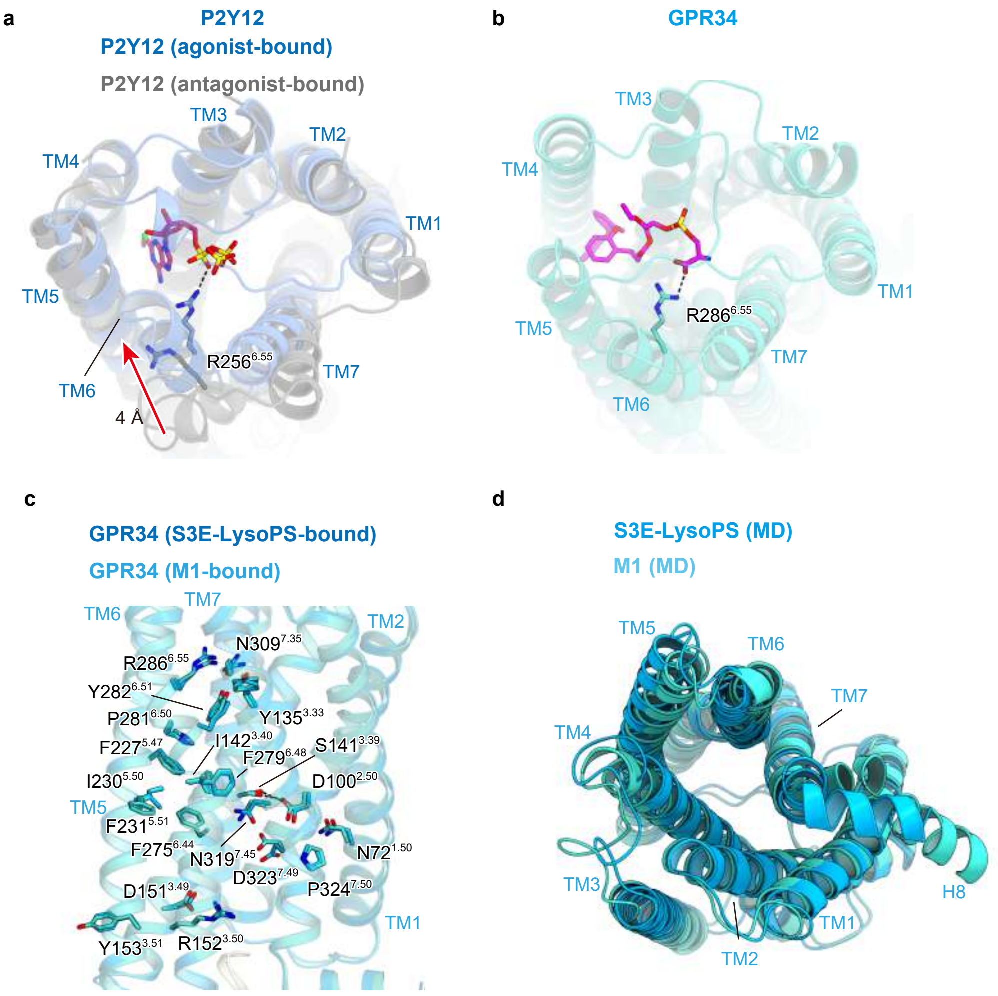


#### Extended Data Fig. 8. Comparison of the conserved motifs.

**(a and b)** Comparison of the in M1-bound GPR34 (a) and P2Y12 (b) structures (PDB 4NTJ and 4PX0). **(c)** Comparison of the conserved motifs at the receptor cores in the M1- and S3E-LysoPS-bound receptors. **(d)** Superimposition of the docking structure of the *sn*-1 derivative and the averaged structure of the M1-bound structure during the 1 μs MD simulation.

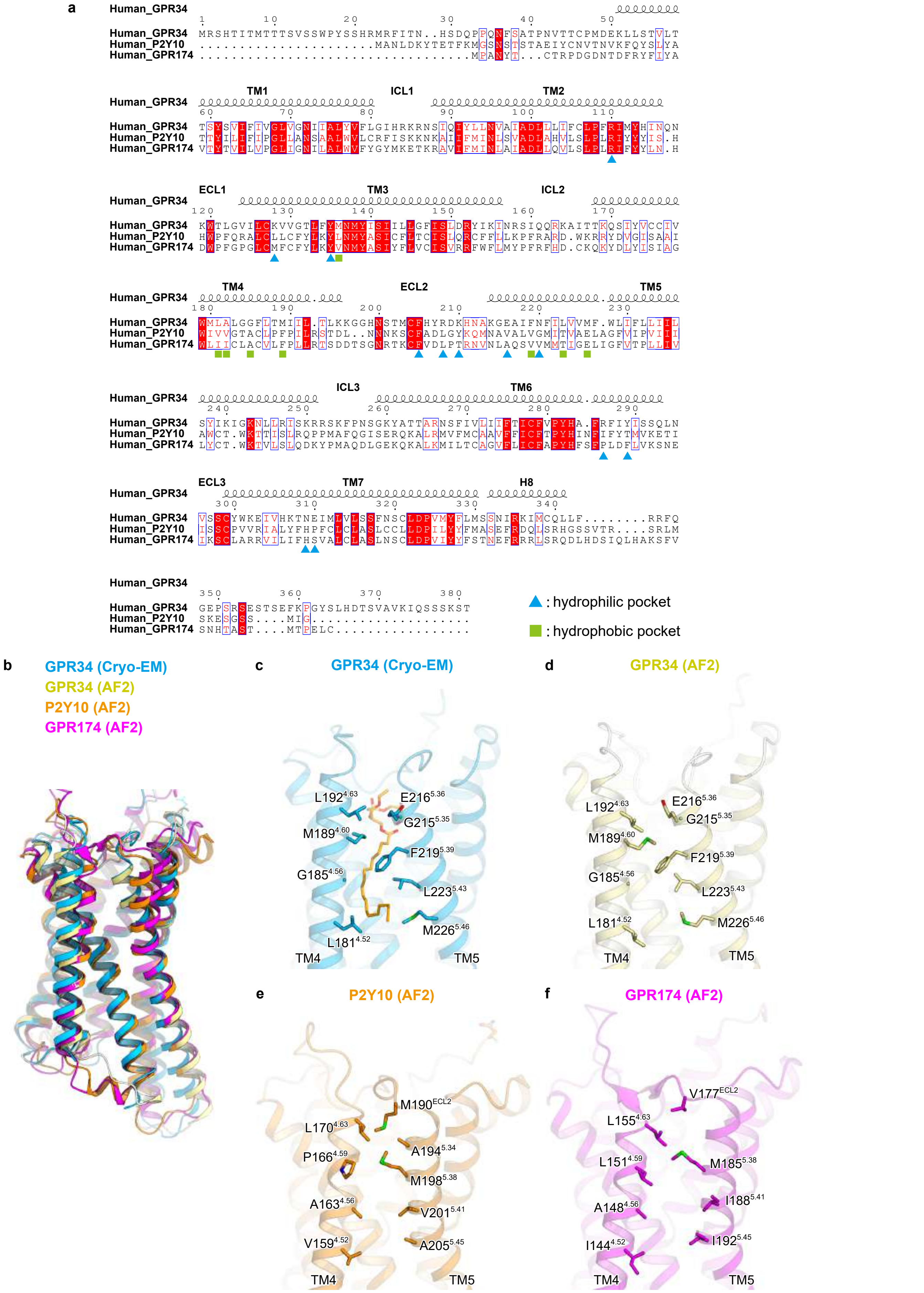


**Extended Data Fig. 9. Comparison of the GPR34, P2Y10, and GPR174 sequences.**

**(a)** Alignment of the amino acid sequences of human GPR34 (Uniprot ID: Q9UPC5), human P2Y10 (Uniprot ID: O00398), and human GPR174 (Uniprot ID: Q9BXC1). Secondary structure elements for α-helices are indicated by cylinders. Conservation of the residues is indicated as follows: red panels for completely conserved, red letters for partially conserved, and black letters for not conserved. **(b)** Superimposition of the structures of GPR34 and AF2 models of human GPR34, P2Y10, and GPR174. **(c-f)** Comparison of the TM4-5 gaps in the cryo-EM GPR34 structure (c) and the AF2 models of human GPR34 (d), P2Y10 (e), and GPR174 (f).
